# Assembly formation is stabilized by Parvalbumin neurons and accelerated by Somatostatin neurons

**DOI:** 10.1101/2021.09.06.459211

**Authors:** Fereshteh Lagzi, Martha Canto Bustos, Anne-Marie Oswald, Brent Doiron

## Abstract

Learning entails preserving the features of the external world in the neuronal representations of the brain, and manifests itself in the form of strengthened interactions between neurons within assemblies. Hebbian synaptic plasticity is thought to be one mechanism by which correlations in spiking promote assembly formation during learning. While spike timing dependent plasticity (STDP) rules for excitatory synapses have been well characterized, inhibitory STDP rules remain incomplete, particularly with respect to sub-classes of inhibitory interneurons. Here, we report that in layer 2/3 of the orbitofrontal cortex of mice, inhibition from parvalbumin (PV) interneurons onto excitatory (E) neurons follows a symmetric STDP function and mediates homeostasis in E-neuron firing rates. However, inhibition from somatostatin (SOM) interneurons follows an asymmetric, Hebbian STDP rule. We incorporate these findings in both large scale simulations and mean-field models to investigate how these differences in plasticity impact network dynamics and assembly formation. We find that plasticity of SOM inhibition builds lateral inhibitory connections and increases competition between assemblies. This is reflected in amplified correlations between neurons within assembly and anti-correlations between assemblies. An additional finding is that the emergence of tuned PV inhibition depends on the interaction between SOM and PV STDP rules. Altogether, we show that incorporation of differential inhibitory STDP rules promotes assembly formation through competition, while enhanced inhibition both within and between assemblies protects new representations from degradation after the training input is removed.

## Introduction

A fundamental link between systems and circuits neuroscience is that the recurrent wiring of neuronal networks is functionally organized [1], and can reflect sensory experience [2] For example, excitatory neurons in the visual cortex show strong reciprocal wiring when neuron pairs have similar stimulus tuning [3, 4, 5, 6], and neurons in the hippocampus form groups of co-activated assemblies that may be used in memory formation [7, 8]. A group of neurons with strong recurrent wiring and shared stimulus or task selectivity is often termed a *neuronal assembly* [9]. A signature of strong assembly structure is that the dynamic and trial-to-trial variability in the joint spiking activity of neuron pairs also reflects their functional structure – neuron pairs within a common assembly show correlated fluctuations in their activity when compared to neuron pairs that are members of distinct assemblies [10, 5, 11, 12]. Thus, tracking how correlated variability across a neuronal population changes with repeated experiences offers a window into the learning mechanisms which carve assembly structure within a network.

Assembly wiring can be learned through an unsupervised Hebbian plasticity rule between excitatory neurons, whereby co-activated neurons form strong reciprocal synaptic connections [13, 11, 12, 14]. However, such strong recurrent excitatory connections can in principle destabilize network activity through unchecked positive feedback. In cortex such network instabilities are alleviated by having strong recurrent excitation dynamically tracked and balanced by an opposing strong recurrent inhibition [15, 16, 17, 18, 19, 20]. In networks that have unstructured wiring such balanced excitation and inhibition promote an asynchronous network state [21, 22, 23, 24, 25]. However, when wiring structure is embedded within an excitatory network, the resulting network dynamics are heavily shaped by how any structure in the inhibitory wiring relates to the excitatory wiring [25, 26, 27, 28, 29, 30]. The specifics of structured inhibition present distinct problems for assembly learning within the excitatory network, as we introduce with a simple motivating example.

Consider a network with a baseline excitatory strength *J* and an embedded pair of assemblies (E1 and E2) where the within assembly excitatory strength is *wJ* (*w* > 1 reflects a Hebbian assembly; Figure 1). We first consider structured recurrent inhibition whereby each excitatory assembly receives strong inhibition from a paired inhibitory assembly (I1 and I2; Figure 1A). In this case as *w* is increased the recurrent inhibition maintains approximate asynchronous spiking dynamics (Figure 1C1 and C2). Thus, while network stability is maintained it now comes at a price – the assembly structure (*w* > 1) is not reflected in the spiking dynamics of the network i.e competition between assemblies is weak. The opposite structure in inhibitory wiring would be lateral connections that mediate competition between the excitatory assemblies (Figure 1B). In this network as *w* is increased a clear competition between the assemblies manisfests (Figure 1D1 and D2). This dynamic certainly reflects the excitatory assembly structure, yet now with the price of pathological behavior when *w* is large, where each assembly either shows high activity with very coordinated spiking or near complete silence (Figure 1D2). In total, this exercise exposes a concrete dilemma. Inhibition can either significantly mask or greatly exaggerate the dynamical signature of embedded excitatory assemblies. This contrasts with the reality that structured excitatory wiring is clearly, yet moderately, reflected in the network dynamics [5, 31, 32]. The central goal of this study is to uncover how real cortical networks use inhibition to ensure network stability yet permit reasonable within assembly correlated activity and between assembly competition.

**Figure 1:**
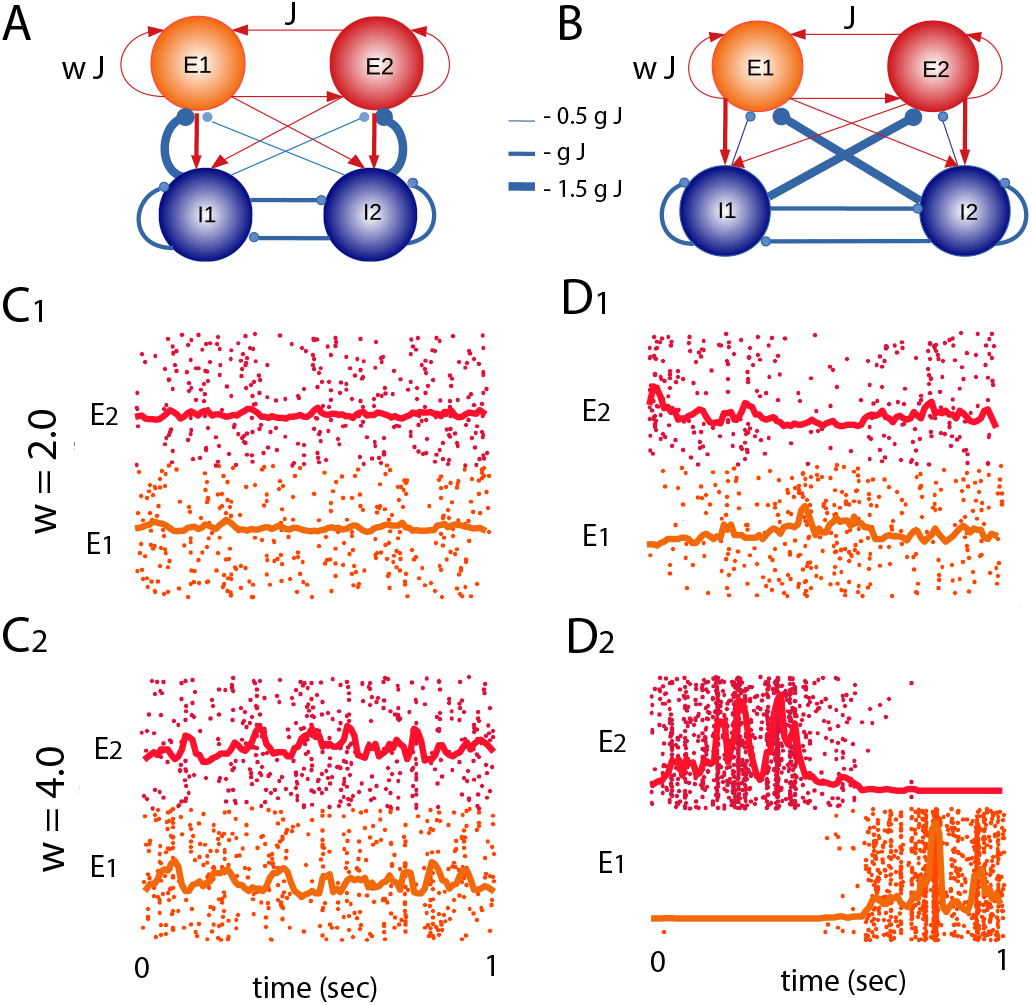
Stability versus competition. **A**: Schematic of a network with bulk inhibition. The scalars *w* and *g* change the relative strength of the weights within the excitatory assemblies, and the projection weights from the inhibitory population onto the excitatory assemblies. **B**: Schematic of a network with lateral inhibition. The inhibitory weights which form lateral inhibition are 3 times as strong as the weights from the inhibitory populations to the excitatory assemblies. **C**_1_, **D**_1_: Raster plots of the networks in A and B for *g* = 6 and *w* = 1. **C**_2_, **D**_2_: Raster plots of the networks in A and B for *g* = 6 and *w* = 3.5. There is a stronger competition in the network with lateral inhibition. **C**_3_, **D**_3_: Raster plots of the networks in A and B for *g* = 6 and *w* = 4.5. The network with lateral inhibition demonstrates episodes of synchronous regular dynamics due to lack of stability in this regime.

It is now well established that inhibition is diverse within most neuronal networks, both in terms of its cellular structure and position within the cortical circuit [33]. Over the past decade a popular hypothesis has emerged whereby this diversity allows distinct inhibitory classes to participate in distinct cortical functions such as providing stability and controlling population-wide correlated activity [34, 35, 36, 37]. In a parallel series of studies inhibition has also been shown to be plastic [38, 39, 40, 41, 42, 43, 44, 45], and whose activity dependent learning mirrors the Hebbian learning in excitatory networks. In our work we explore the hypothesis that cell specific inhibitory plasticity onto excitatory neurons allows a diverse inhibitory network to simultaneously confer both stability and assembly based network competition. In an *in vitro* preparation of mouse orbital frontal cortex (OFC) we use a spike timing dependent plasticity (STDP) protocol and uncover that parvalbumin-expressing (PV) interneurons have a homeostatic and symmetric STDP learning rule, while somatosatin-expressing (SOM) interneurons have a causal Hebbian learning STDP rule. Armed with these inhibitory STDP rules we use a combination of simulations of a model cortical network and an associate mean-field theory to show how a cortical network with PV and SOM neurons support the rapid training of stable excitatory assemblies whose structure is reflected in the ongoing spiking dynamics. In particular, the different inhibitory plasticity rules result in a division of labor: the PV network confers a homeostatic stability of the network dynamics (Figure 1A), thereby permitting the SOM inhbitory network to support cross assembly competition (Figure 1B), however, SOM populations provide competition between excitatory assemblies as a result of the formation of lateral inhibition. In addition to this, we also show that SOM neurons could also potentially result in a subtle emergence of tuned PV to excitatory connections, by correlating PV and excitatory neurons given the lateral connections to the excitatory subnetwork and heterogeneous connections to PV neurons. Our work thus provides an important mechanistic insight into how diverse inhibitory learning supports the stable creation of learned excitatory networks.

## Results

### Experimental findings

Previous studies have established the importance of the temporal window between synaptic input and evoked action potentials in the modification of excitatory synaptic strength[46]. However, fewer studies have investigated the STDP of inhibitory synapses[43, 39, 47] particularly between specific subclasses of interneurons and excitatory pyramidal neurons (PNs). We expressed channelrhodopsin (ChR2) in PV or SOM interneurons then investigated the STDP of light-evoked inhibition onto PNs in L2/3 of orbitofrontal cortex (OFC). Briefly, STDP was induced by pairing light-evoked inhibitory post synaptic potientials (IPSPs) with evoked action potentials in the PN (See Methods, Figure 2A). For each recorded neuron, inhibition was evoked at a single temporal interval defined as the difference between the time of the AP (t_spike_) and the time of the IPSP (t_IPSP_) (Figure 2A1). Negative interval values correspond to AP’s arriving prior to the IPSP, while positive values correspond to APs following the IPSP (Figure 2A2). To assess the time course of synaptic plasticity, baseline IPSCs (Figure 2B1, C1) were collected in voltage clamp for 5 min prior to the STDP pairing protocol and test IPSCs were collected for up to 1 hour following (Figure 2B2,C2). Synaptic plasticity was quantified as the average test IPSC amplitude 25-30 min post induction (gray shading, Figure 2B2,C2) normalized to baseline IPSC amplitude.

**Figure 2:**
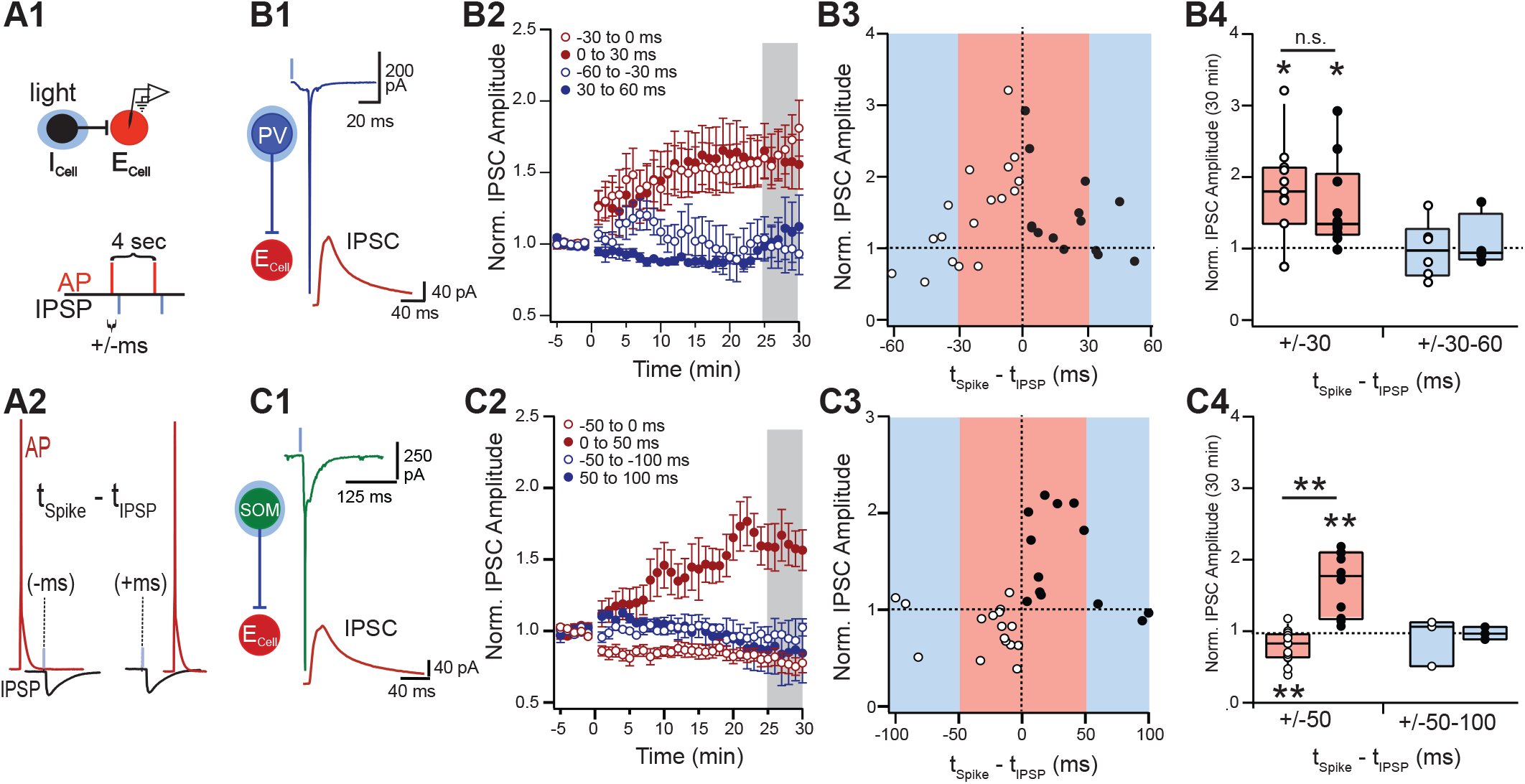
Inhibitory STDP differs between PV and SOM interneurons **A**: Schematic of STDP pairing protocol. Action potentials (AP) are evoked in the post-synaptic pyramidal neurons(E_Cell_) every 4 seconds (0.25 Hz) (a1,a2 red) while light evoked IPSPs are evoked at a constant interval (ms) relative to the AP (a2,black). **B**: Inhibitory STDP from PV interneurons is symmetric. B1: Light evokes a single AP in a ChR2+ presynaptic PV interneuron (blue trace). IPSCs (red trace) are recorded in the post-synaptic E_Cell_. B2: Baseline IPSCs are collected 5 min prior to the STDP protocol (time =0) and up to 60 min post pairing. IPSC amplitude is normalized to the average baseline amplitude. B3: Normalized IPSC amplitude recorded 25-30 min post STDP induction (B2, gray shaded) plotted against temporal window for STDP pairing (ms). Negative intervals correspond to post-synaptic spikes before the IPSP (open circles), positive intervals correspond to IPSPs before the spike (closed circles). B4: Normalized amplitude was binned in 30 ms intervals for comparison between positive and negative intervals. Intervals between 0 and +/-30m ms show significant potentiation,*p<0.05, but do not significantly differ between positive and negative intervals. **C**: Inhibitory STDP for SOM interneurons is asymmetric. C1-C3: As for b1-b3. C4) Normalized amplitude was binned in 50 ms intervals for comparison between positive and negative intervals. Short positive intervals 0-50 ms show significant potentiation (**p<0.01) whereas short negative intervals -50-0 ms show significant depression (**p<0.01). Short intervals significantly differ between positive and negative intervals.

We find that inhibitory STDP rules differ between PV and SOM interneurons. Inhibition mediated by PV interneurons is significantly potentiated for short intervals within +/-30 ms of the post-synaptic action potential (red shaded regions, Figure 2B3) and declines as intervals increase from +/-30-60 ms (blue shaded regions). We binned normalized amplitude in 30 ms bins and found that synaptic potentiation was significantly >1 for short-intervals (−30-0 ms: 1.79 +/-0.69, p=0.004, n=11; 0-30 ms: 1.60 +/-0.62, p=0.010, n=10, single distribution t-test, Figure 2B4). This relationship was symmetric with respect to the timing of IPSPs pre versus post the AP as synaptic enhancement did not significantly differ for positive versus negative intervals (p=0.532, unpaired t-test; Figure 2B4). Conversely, the temporal dependence for STDP of SOM inhibition was asymmetric. Inhibition was significantly potentiated (Norm. Amp. >1) for positive intervals between 0-50 ms when inhibition preceded spiking (1.66 +/-0.44, p=0.001, n=10, Figure 2C3,C4). Whereas inhibition was significantly depressed (Norm. Amp. <1) for negative intervals (−50-0 ms: 0.78 +/-0.22, p=0.004, n=13). Synaptic plasticity significantly differed between positive and negative intervals <50 ms (p=1.5E-5, unpaired t-test, Figure 2C4).

### Correlated population spiking activity induced by SOM and PV plasticity

To explore the emerging network properties as a result of PV and SOM plasticity, we include the learning rules of these inhibitory subtypes in networks of model neurons with spiking dynamics (leaky integrate-and-fire; see Methods). We model assembly formation by separating the excitatory neuron (E) into two subnetworks (E1 and E2; Figure 3A-C) based on their exposure to a fluctuating training signal that will interact with a Hebbian E → E STDP plasticity to form E assembly structure [11, 13, 48]. In what follows there is no initial E → E structure (i.e. E1 and E2 have identical within and between connectivity strengths), and any final structure must emerge through assembly training.

**Figure 3:**
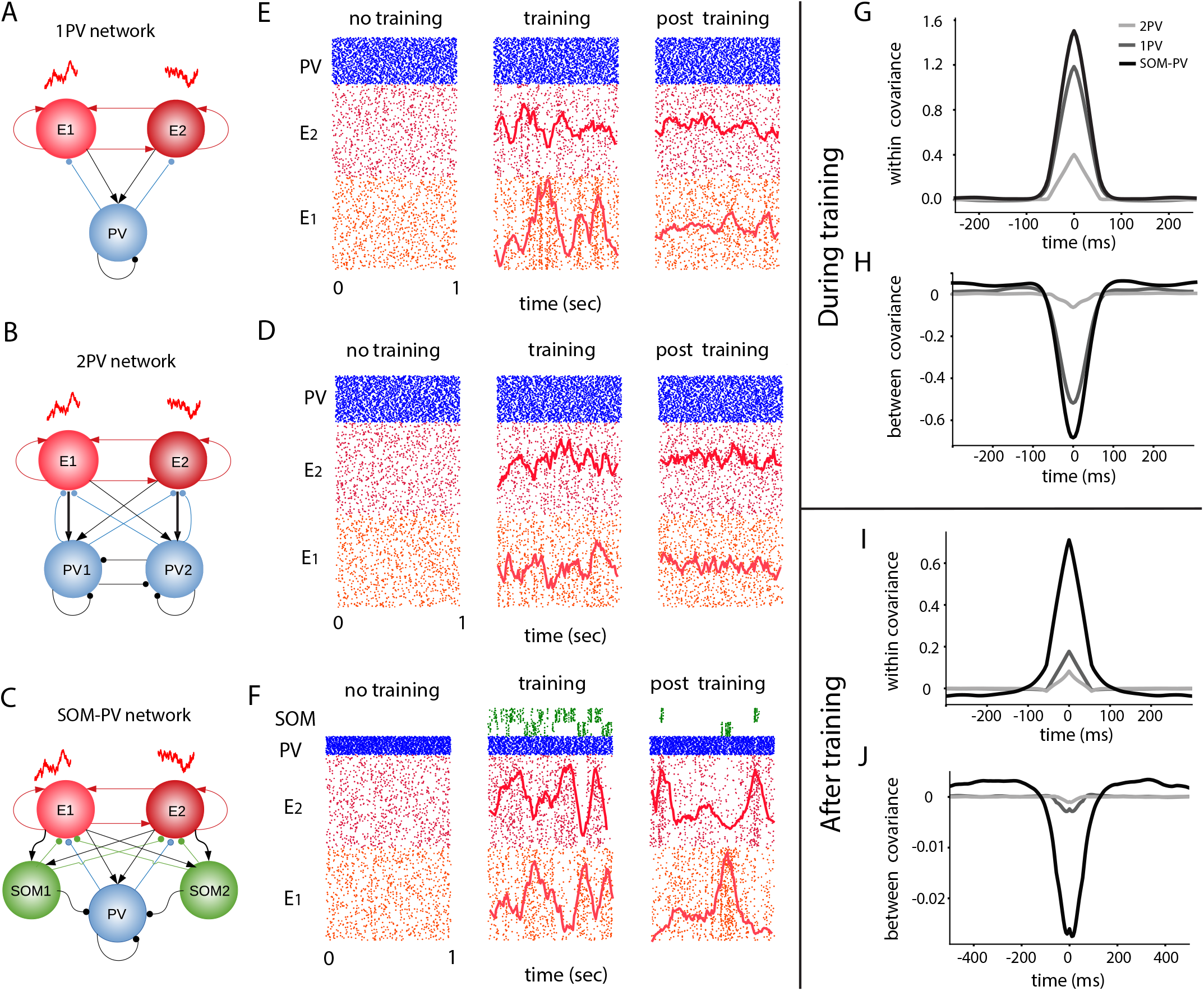
Different network architectures and the resulting correlation structure during and after training. **A**: 1PV network has 2 separate groups of excitatory neurons, and one shared group of PV cells. There are no tuned connections in this case. **B**: 2pv network is composed of 2 distinct groups of excitatory neurons which are tuned to distinct groups of PV neurons. **C**: SOM-PV network is similar to the 1pv network, but in addition, it has 2 groups of distinct SOM neurons, each of which receive tuned connections from excitatory neurons. **D, E, F**: raster plosts of the networks in panels A, B, C, respectively, when there is no training, during training, and after training. Only 3 seconds of the network activity has been shown for each case. **G, I**: Autocorrelation function for the average population firing rate of a single excitatory assembly during training, and after training, respectively, for each network architecture. **H, J**: Cross-correlation function between the average firing rates of each excitatory assembly during and after training, respectively.

We consider three different network models, distinguished by their inhibitory subnetworks. The first is a ‘1PV network’ (Figure 3A) with two excitatory populations and one global inhibitory population, without any tuned connections. The second is a ‘2PV network’ (Figure 3B), composed of E and PV cells where neurons in the E1 population connect more strongly and with a higher probability to half of the PV cells in the PV population (the E2 population connects more strongly to the other half). This network is composed of the same number of PV neurons as the 1PV network. Finally, the ‘SOM-PV’ network (Figure 3C) has two different inhibitory cell types, PV and SOM class interneurons. The connections from E to PV are not structured, but we assume a tuned connection from E to SOM cells, such that half of the excitatory neurons are more strongly and with a higher probability are connected to a subset of SOM population (similar to the 2PV network). In all cases the E → E, PV → E, and SOM → E synaptic connections have the appropriate STDP learning rules (see Methods; colored connections in Figure 3A-C denote plastic connections).

Before training, all networks express asynchronous, temporally irregular spiking activity (Figure 3E-F). SOM neurons in the SOM-PV network received synaptic inputs only from the E population, along with an external static current to bring up the resting membrane potentials in SOM neurons; nevertheless, before training, SOM neurons have negligible firing rates (Figure 3F). During training, each E neuron receives external excitatory synaptic trains with Poisson interval statistics. There are two sources of inputs to each neuron. The major inputs are unshared sources that feed each neuron independently with 70 % of the total external input firing rate. In addition to this unshared input, there are also two independent sources of Poisson processes, each of which provide a shared input (the remaining 30 % of the total external input) to two individual subgroups of the excitatory population. The goal of the shared input is to correlate neurons in each excitatory sub-population independently of the other population. The shared source could for example represent a different odor which acts as a different sensory stimulus. During training, neurons in each excitatory population that receive shared input are highly correlated. This correlation manifests itself in the correlation function between average neuron spontaneous firing rates which reside in each population (the average PSTH function for each excitatory population is shown on the raster plot for reference). The amplitude of the auto-correlation function for the average firing rate in each assembly is maximum for the SOM-PV network, and minimum for the 2PV network (Figure 3G). Moreover, the cross-correlation function between the average firing rates of the two excitatory populations is more negative for the SOM-PV network than the cross-correlations for other networks under study (Figure 3H). This implies that competition between cell assemblies during training is stronger in the SOM-PV network. Moreover, the bursty nature of activity of SOM neurons in the SOM-PV network is due to lack of connections between SOM neurons and the fact that they only receive excitation from the activity of the excitatory populations. An experimental observation reporting a similar phenomenon can be found in [49].

After 2000 seconds of training, we removed the shared source and replaced it with a global independent unshared stimulus source. It is noticeable that the activity of excitatory neurons in each assembly remains highly correlated in the SOM-PV network, and this correlation structure is much less impressive in the 1PV and 2PV network (Figure 3I). Similar to the competition structure between cell assemblies during training, the cross-correlation between assembly firing rates after training is more impressive in the SOM-PV network (Figure 3J). This hints to the fact that after training, the assembly structure in the SOM-PV network has maintained itself to a larger degree. Overall, we conclude that the existence of SOM neurons with Hebbian STDP, during and after training, has resulted in a bigger amplitude of auto and cross correlation function between average neuronal firing rates in each assembly.

### Excitatory weight evolution in different networks

To explain how the connection weight between two neurons changes, we consider a pre-synaptic neuron *j* impinging on a postsynaptic neuron *i* (Figure 4A). Each neuron fires according to a pattern. In order to calculate the average weight evolution between the pre-synaptic and a post-synaptic neuron, it is conceivable to first calculate the cross-correlation function between the spike trains of the pre-synaptic and the post-synaptic neuron (Figure 4B). The integral of the product of this function with the STDP curve will yield the average rate of change in the connection weight between the two neurons [50]. This integral, after being scaled by a learning rate *η* is the sum of all average potentiation (indicated by *U* 1 in Figure 4C) and depression (indicated by *U* 2 in Figure 4C) that the synapse undergoes. We use this idea throughout the paper to justify the average weight change between populations of neurons. For this, we assume that the average spontaneous firing rate of neurons in each population stands for the firing rate of individual neurons in a mean-field sense.

**Figure 4:**
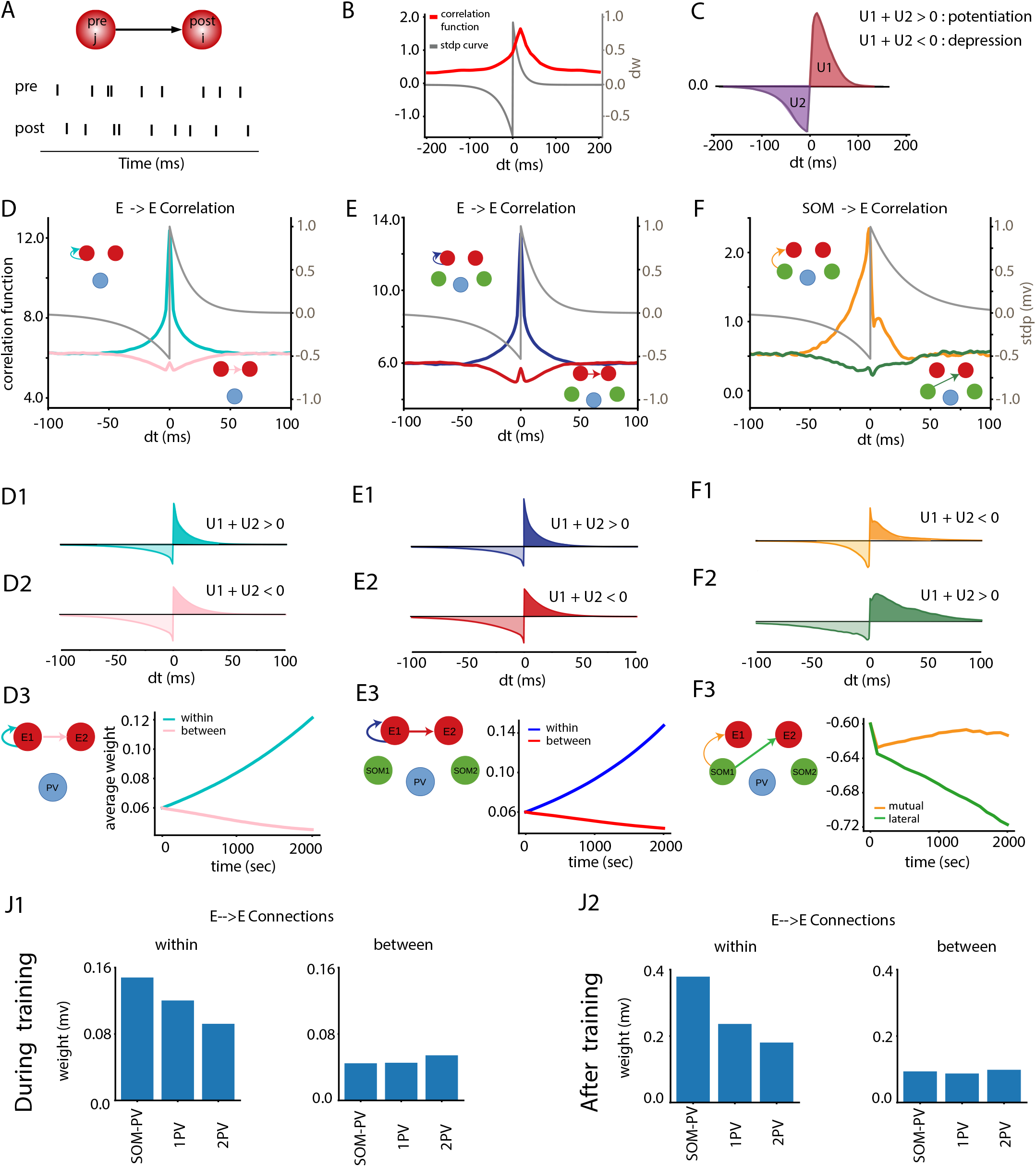
Correlation functions and weigh evolution. **A**: Spiking activities of a presynaptic neuron *j* which projects onto a postsynaptic neuron *i*. On average, the presynaptic neuron fires before the postsynaptic neuron. **B**: STDP curve is plotted in gray, and the correlation function between the spike trains of the presynaptic neuron with the postsynaptic neuron is plotted in red. **C**: The integral of the cross-correlation function and the STDP curve results in a scalar with a positive part (*U*_1_), and a negative part (*U*_2_). The sign of the sum of these two quantities determines whether the synaps potentiates or depresses. **D, E, G**: Within assembly autocorrelation function (solid lines) and between assebmly cross-correlation function (dashed lines) for the 1pv network, SOM-PV network, and 2PV netwrok, respectively. The Hebbian STDP curve which rules the weight update between excitatory neuron is plotted in gray. **D1, D2**: The integral of the product of the STDP curve with the single assembly auto-correlation function and the cross-correlation function between assemblies in panel D, respectively. **D3**: Within assembly (cyan) and between assembly (pink) weight changes as a function of training time. At *t* = 2000 s, the average weights within an excitatory assembly is slightly above 0.12 mV. **E1, E2**: The integral of the product of the STDP curve with the auto-correlation function and the cross-correlation function in panel E, respectively. **E3**: The excitatory weight evolution for the connections within (blue) and between (red) assemblies. At the end of the training time (*t* = 2000 s), the average weight within an excitatory assembly is slightly above 0.14 mv. **F**: Cross-Correlation function beween SOM1 and E1 in solid lines, and cross-correlation between SOM1 and E2 in dashed lines. **F1, F2**: The integral of the product of the STDP curve with the cross-correlation function between SOM1 and E1 (F1), and the cross-correlation function between SOM1 and E2 (F2) in panel E, respectively. **F3**: Development of the lateral and mutual inhibitory weights from SOM1 to E2 and to E1, during training, plotted in green and yellow respectively. The lateral inhibition is stronger than the mutual inhibition. **J1**: During training, at t=2000 sec, the weights within assembly are the largest, and the weights between assemblies are the weakest for the SOM-PV network. **J2**: After 3800 seconds post training, the weights within the excitatory assemblies are the largest for the SOM-PV network, and the smallest for the 2PV network. The weights between assemblies is bigger in the 2PV network.

During training, the average auto-correlation function for the excitatory populations in all networks under study has a peak for zero lag time shifts (Figures 4D,E, 5G). The integral of the product of these functions with the STDP curve for the excitatory connections results in positive scalars, which indicates potentiation of the synapse. This product has been illustrated in Figures 4D1, E1, 5G1 for the 1pv, SOM-PV and the 2PV network, respectively. The area under the curves are positive in darker shades, and negative in lighter shades for each corresponding synapse. The overall integral, the sum of the positive and negative areas, are 1.18 ×10^−5^, 1.41 ×10^−5^, and 7.45 ×10^−6^ mV/s, for the 1pv, SOM-PV and 2PV network, respectively (These results were obtained by considering neuronal average PSTH signals from *t* = 200 to *t* = 700 seconds during training). The auto-correlation function for the SOM-PV network grows in amplitude for progressive points in time, and this growth in amplitude also slightly changes the growth rate of the weight through a positive feedback loop. At *t* = 2000 seconds during training, the average weights within an excitatory assembly is slightly above 0.12 mV for the 1pv network (Figure 4D3). This quantity reaches 0.146 mV for the SOM-PV network, and 0.095 mV for the 2PV network (Figures 4E3, 5H, respectively).

**Figure 5:**
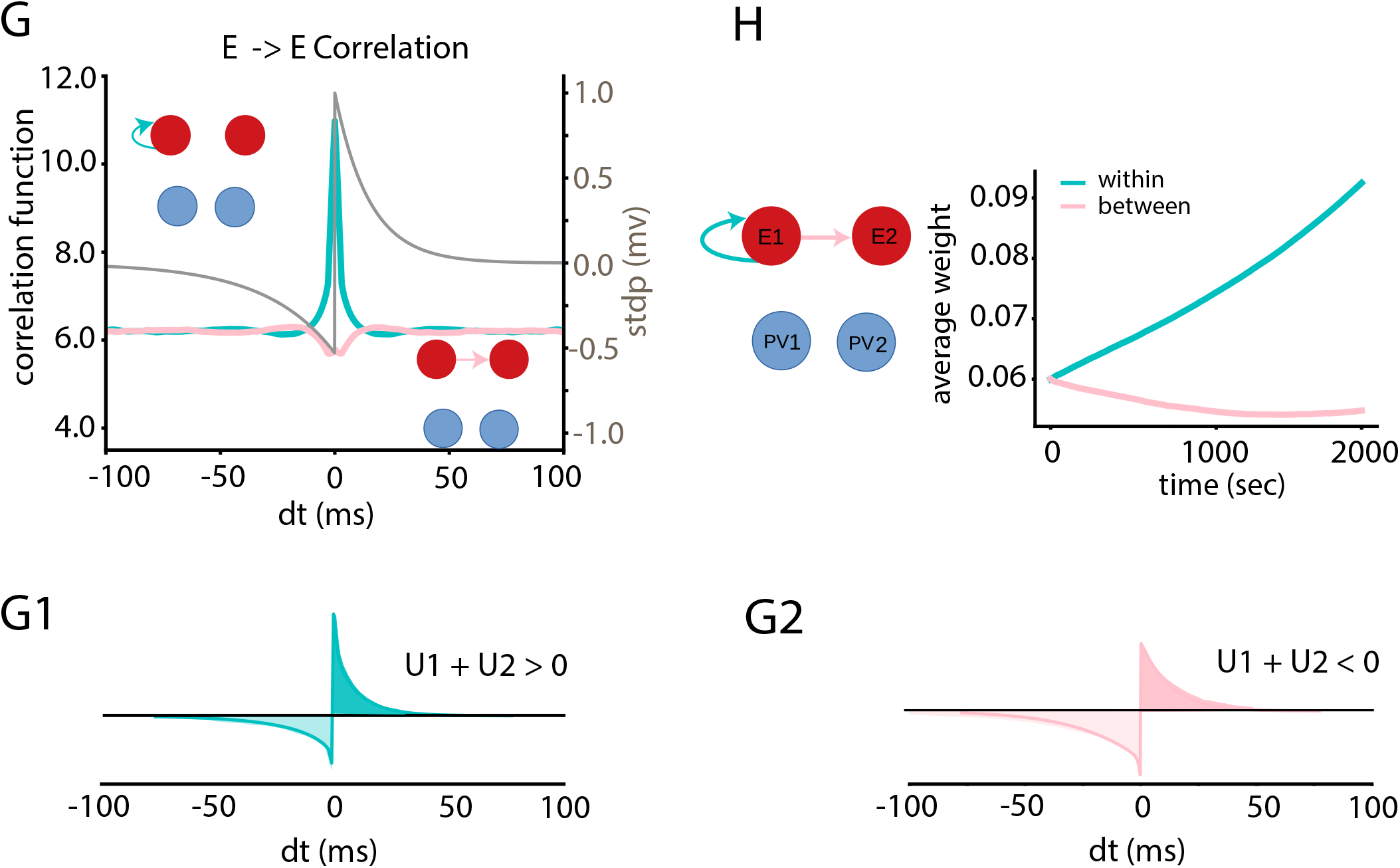
Correlation functions and weigh evolution for the 2PV network. **G1, G2**: The integral of the product of the STDP curve and the correlations in panel G. **H**: Excitatory weight evolutions for the connections within (cyan) and between (pink) assemblies, for the 2PV network.

However, the integral of the product of the STDP curve and the cross-correlation functions between the average firing rates of neurons in different assemblies is negative. This quantity is equal to −3.33 ×10^−6^ mV/s for the 1pv network (Figure 4D2), and is equal to −3.04 ×10^−6^ and −2.21 ×10^−6^ mV/s for the SOM-PV network (Figure 4E2) and the 2PV network (Figures 5G2), respectively. This results in the depression of the coupling weights between neurons in different assemblies (Figures 4D3, E3, H, pink and red curves). This depression of the average weights between assemblies improves assembly segregation, and facilitates distinct assembly formations corresponding to different input signals.

### Role of PV homeostasis in the evolution of excitatory weights

The STDP curves for the excitatory connections had a net negative integral. This indicates that if there was no correlation structure between the pre-synaptic and post-synaptic activity of neurons, the average synapse between those neurons would depress. The fact that the assemblies are being formed due to the potentiation of the synapses within assemblies is because the product between the “cross-covariance” function between neurons and the STDP curve is a positive scalar and bigger than the product of the mean firing rates and the STDP curve (In this argument we rely on the fact that a cross-correlation function is the sum of the cross-covariance function and the product of the mean firing rates of individual neurons). This picture is different for the synapses between neuron in two distinct assemblies. Due to competition between neurons in different assemblies, the cross-covariance between those neurons is mostly negative. Hence, the total integral of the product of the STDP curve and the correlation function is negative, and as a result, the synapses between assemblies decay with time. In all networks under study, the homeostasis provided by the connections from PV to E cells guaranteed that the firing rate of the excitatory neurons stay fixed and close to 3 Hz. Therefore, the difference in growth rate of the excitatory synapses in the networks under study is mainly due to the different cross-covariance structure between the activities of the excitatory neurons. We will show in the rest of the paper that this covariance is shaped by the SOM plasticity in favor of speeding up the growth of the synapses between neurons in excitatory assemblies.

### SOM plasticity results in lateral inhibition

The tuned connections from each excitatory assembly to their corresponding SOM population results in a causal activity of SOM after its corresponding excitatory activity visits the high state of its spontaneous fluctuations. This causality, due to the Hebbian nature of the plasticity, results in a left-skewed correlation function between SOM_1_ and E_1_, as shown in Figure 4F. As a result, the integral of the superposition of this function on the STDP curve will be a negative quantity (Figure 4F1, −2.39 ×10^−7^ mV/s for correlations between signals that represent average neuronal activities from *t* = 200 to *t* = 300 seconds), which implies the depression of the synaptic weight between SOM_1_ and E_1_ (and similarly the weight between SOM_2_ and E_2_). On the other hand, the activity of neurons in SOM_1_ suppresses the neurons in E_2_. This results in a trough for positive and small time lags in the cross-correlation function (Figure 4F). However, since the integral of the STDP curve is positive (Figure 4F2, 6.2 × 10^−6^ mV/s for correlations between signals that represent average neuronal activities from *t* = 200 to *t* = 300 seconds), the overall weight change would result in a positive quantity and consequently the potentiation of the synapse. Over time, this potentiation forms lateral inhibition from the SOM population that is tuned to the opponent excitatory assembly onto the other excitatory assembly which is suppressed as a result of the activity of those SOM cells. Subsequently, the lateral inhibition amplifies the negative cross-correlation between the excitatory populations, and hence stronger competition between assemblies. Moreover, as a result of this competition, single population auto-correlation functions also increase in magnitude over time. As inferred from [50], this increase in auto-correlation function causes faster formation of cell assemblies. Also, the competition between assemblies which reflects in the cross-correlation function between the excitatory assemblies results in a weak coupling between the two assemblies.

In summary, the emergence of lateral inhibition shapes the correlation functions in such a way that both the auto- and the cross-correlation functions increase in amplitude over time, and this facilitates and speeds up assembly formation. We compared the weights within and between the excitatory assemblies during training after 2000 seconds. The weights between assemblies were the minimum, and the weights within an assembly were the maximum for the SOM-PV network (Figure 4J1). This means that the 1pv and the 2PV network, in comparison with the SOM-PV network, show a slower formation of assemblies. Also, the least segregation between assemblies was observed in the 2PV network. Moreover, 3800 seconds after training, we checked those weights and observed that all weights had potentiated. This potentiation was stronger for the weights within assemblies (Figure 4J2). This is because in all these networks 2000 seconds of training was enough time to form assemblies with correlation functions that were big enough to maintain assemblies (see [11] for more details). The structure of the correlations were such that the assembly structure was the best preserved and grown in the SOM-PV network, and the growth was the slowest for the 2PV network. In the rest of the paper, SOM-PV and 1PV networks will be compared with each other. We will refer to the 2PV network later in the Results section.

### SOM facilitates assembly formation and slows down assembly deformation

For more realistic parameters of the STDP curve (integral of the total STDP curve comparable to the integral of the curve for positive lag) and slightly lower input firing rates, we simulated the 1pv and SOM-PV network for different duration of the training time. Since the STDP curve is depression dominant, we did not provide a training signal to the network for 1000 seconds until all transient dynamics in the weight evolution brought the excitatory weights to their minimum, and similar initial conditions for both 1pv and SOM-PV network. For similar initial conditions in the excitatory weights for both networks, a training signal was provided to both networks. After 1000 seconds of training, the SOM-PV network developed a slightly stronger assembly structure compared to the 1pv network (Figure 6A). At *t* = 0, the training signal was removed. Right after training, the weights within assemblies started to decay, and there was a slight potentiation in the weights between assemblies. The depression for the 1pv network was stronger and happened faster for the 1pv network. The SOM-PV network, however, regained the assembly structure about 1500 seconds after training. In a separate simulation, we increased the input training time to 4000 seconds, and observed that assemblies were formed much faster in the SOM-PV network (Figure 6B). After the training signal was removed at *t* = 0, assemblies continued growing after a slight deformation. This growth can be attributed to the remaining correlation structure which can maintain itself and get amplified even when the training signal does not exist. A similar phenomenon was reported in [51, 11], where internally generated correlations as a result of training were sufficient to drive the weight evolution even after training. At *t* = 5000 s post training, the weights within individual assemblies are much bigger for the SOM-PV network.

**Figure 6:**
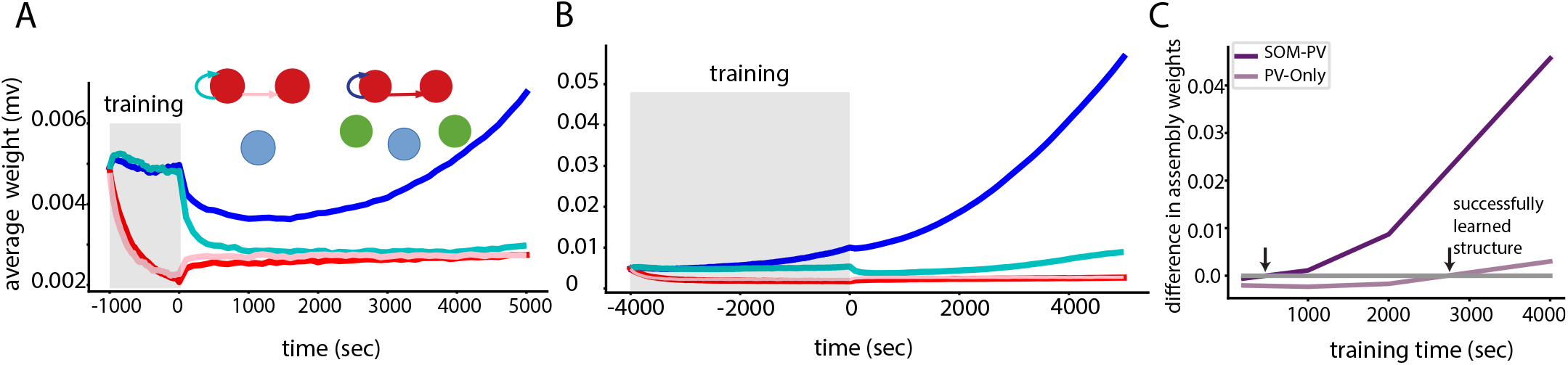
Faster training and slower forgetting in the SOM-PV network. **A**: For a training period of 1000 seconds, the excitatory weights within each assembly grow faster in the SOM-PV network compared to the 1pv network. After training, the weights within and between assemblies are almost identical in the 1pv network. **B**: For a prolonged training period of 4000 seconds, assemblies continue to grow their structure after training. The growth rate is faster in the SOM-PV network. **C**: 4000 seconds after the training signal was removed (*t* = 4000 s), the difference between the weights within each assembly and between assemblies (*dw*_*t*_) were compared to that quantity right after training was stopped (*dw*_0_ at t=0 s). The index *t* in *dw*_*t*_ represents the training time. When the difference crosses 0, it means the same assembly structure right after training is recovered at *t* = 4000 s. Zero-level crossing happens in less training time for the SOM-PV network.

In order to quantify the training time necessary to maintain the assembly structure similar to the existing one at *t* = 0 post training, we referred to a measure which was the difference between the weights within and between assemblies, called dw_0_. We also calculated this measure for *t* = 4000 for different network simulations with different training times and called this dw_4000_. Then, we plotted dw_4000_ - dw_0_ for different training times for the SOM-PV and the 1pv network (Figure 6C). The training time for which this measure crosses zero is the minimum training time necessary to recover the same assembly structure at *t* = 4000 s that existed right after training (*t* = 0). As our results suggest, this training time is much smaller for the SOM-PV network compared to the 1pv network. This concludes that it is easier to train and maintain an assembly structure when SOM inhibitory plasticity contributes to assembly formation.

To understand the dynamics of the excitatory weights during assembly formation, we considered a dynamical system analysis in the weight state space. To characterize assembly formation, we were interested in two states: weights within assemblies (*w*_11_) and weights between assemblies (*w*_12_). We conducted a mean-field analysis for the average neuron’s correlation functions within individual populations. These correlations are functions of the plastic and static weights. Given that the SOM-PV network has 5 interacting populations (*E*_1_, *E*_2_, *I, SOM*_1_, and *SOM*_2_), and 5 plastic weights (*w*_11_, *w*_12_, *w*_*i*_, *w*_*s*1_, and *w*_*s*2_), the entire set of coupled systems of ODEs is 10 dimensional (see the Methods section for details). A similar analysis for the 1pv network results in a 6 dimensional system. In order to get a phase plane analysis in the parameter space of interest (*w*_11_ and *w*_12_), we used the Laplace transform to relate the dynamics of the weights to the firing rates, and performed a dimensionality reduction in the Laplace domain such that the entire system could be described as a function of *w*_11_ and *w*_12_ only.

The phase plane analysis for the dynamics of *w*_11_ and *w*_12_ for the 1pv and the SOM-PV network suggests that in a given parameter range close to the origin, the nullcline associated with *w*_11_ is closer to the origin for the SOM-PV network (Figure 7A,B). This means that the area in the parameter space in which learning takes place (arrows that point to the right and downwards) is bigger for the SOM-PV network. Also, the velocity of the vectors associated to the *w*_11_ component is bigger for the SOM-PV network compared to the 1pv network. This automatically implies that training assemblies is faster in the SOM-PV network.

**Figure 7:**
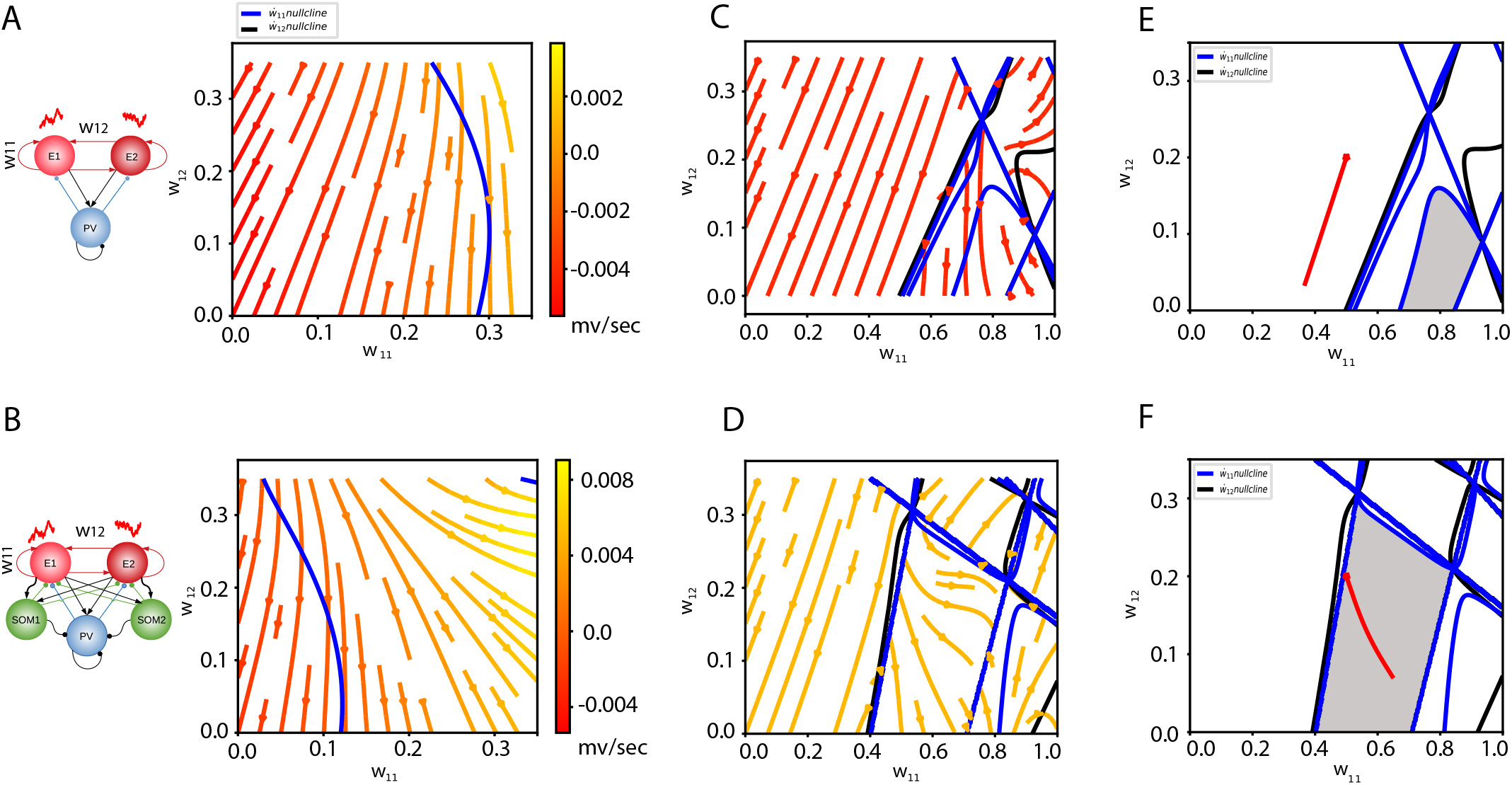
Mean-field phase portraits for the excitatory weight evolution for the two networks. **A**: When the training signal is present (c=0.3), the nullcline for the 1pv network is farther from the origin than the nullcline for the SOM-V network. The flow is bigger in the SOM-PV network case. **B**: After training, the nullcline for *c* = 0 is closer to the origin in the case of SOM-PV network. **C**: The region in the parameter space in which assembly growth continues after training is highlighted in gray. For an initial condition of (*w*_11_, *w*_12_) = (0.5, 0.2), which could result from a training phase, the trajectories after the training signal was removed have been plotted in red.

The phase portraits for both networks after training suggest that if *w*_11_ and *w*_12_ are both small, the flow in both directions decreases until the weights converge to zero. However, if the state of the weighs within and between assemblies are such that the nullcline for *w*_11_ is passed, then the flow can continue in the direction of increasing *w*_11_ (Figures 7C,D). This means that the assembly can sustain itself, only if the training time was sufficient enough and as a result the initial conditions post-training were left in the region of interest where assemblies could potentially grow. This region of interest is highlighted in gray in Figure 7E,F for both networks. For an exemplary initial condition (*w*_11_, *w*_12_) = (0.5,0.2), right after training, the flow will move towards the origin for the 1pv case, however, for the SOM-PV network, the flow continues in the direction of increasing *w*_11_ and decreasing *w*_12_. This is all due to the leftward shift of the nullclines for the SOM-PV case which suggests maintaining the assembly structure after training is relatively easy if SOM plasticity contributes to learning. Also, the region of interest is relatively bigger in this case.

### Emergent properties of interaction between inhibitory subtype plasticity

Thus far, we have discussed the connection weights within and between excitatory assemblies in the 3 networks we considered. We also discussed the emergence of lateral inhibition from SOM to E cells in the SOM-PV network. In this section, we investigate the evolution of the inhibitory weights from PV onto E cells.

For all networks, we summed up all the weights that any individual PV neuron impinged on the E cells in E_1_, and called this sum *PV* → *E*_1_. Similarly, we summed up all the projecting weights from any PV cell onto all neurons in E_2_, and called this quantity *PV* → *E*_2_. We then sorted the index of all PV cells such that the neurons with lower indices had the largest *PV* → *E*_1_. The connectivity matrix from the PV cells to the entire E cells for the 1pv network is shown in Figure 8A. The first left block of this matrix is sorted as explained before. The second half on the right does not follow any specific pattern, and therefore, we conclude that for this network, there is no systematic preferential bias for PV cells to connect more strongly to E cells in either of the assemblies.

**Figure 8:**
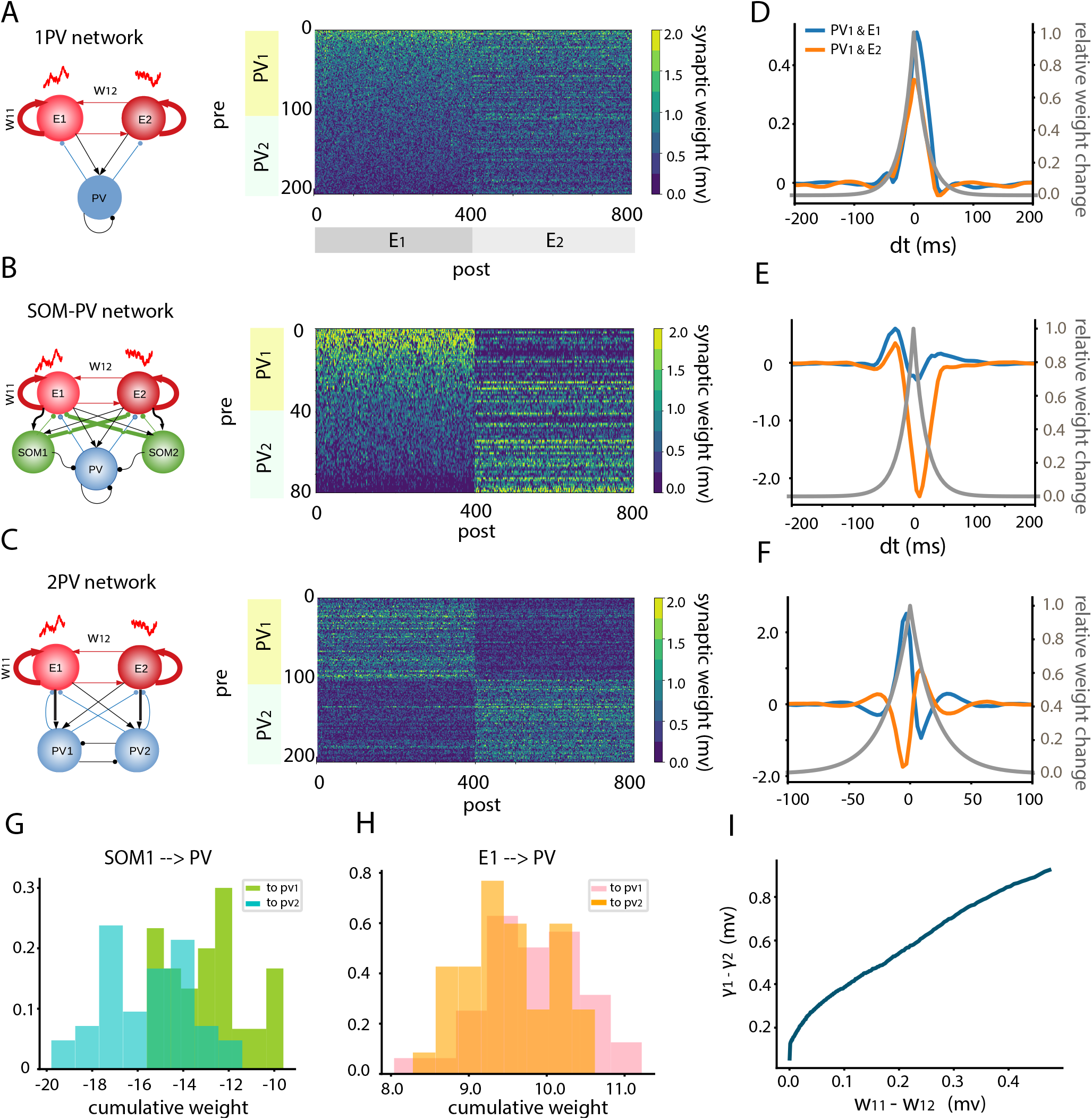
Emergence of tuned PV to E connections in the existence of tuned excitatory connections. **A**: Connectivity matrix from presynaptic PV neurons to postsynaptic E neurons for the 1pv network and the SOM-PV network, respectively. The index of the PV neurons were sorted such that lower indexed neurons project strongly onto neurons in E_1_. After this sorting, the first half of the PV neurons were called PV_1_, and the second half PV_2_. **B**: Connectivity matrix representing the PV to E connections. After sorting the indices of the PV neurons, it appears that PV cells mostly tend to connect strongly either to E1 or E2, but not both. **C**: PV to E connectivity matrix for the 2PV network without sorting any index. **D, E, F**: Cross-covariance function between the average spontaneous firing rates of PV_1_ and E_1_, and also between PV_1_ and E_2_ for the networks represented on the left side of each figure. The symmetric STDP curve for the connections from PV to E neurons is shown in gray. The right axis represents the relative scales of the symmetric STDP curve for the connections between PV and E cells. **G**: Histograms of the cumulative weight projections from SOM_1_ onto PV_1_ and PV_2_ neurons in the SOM-PV network. **H**: Histograms of the cumulative weight projections from E_1_ onto PV_1_ and PV_2_ neurons in the SOM-PV network. **I**: The difference between tuned and untuned PV to E weights scales almost linearly and positively with the difference between the weights “within” and “between” assemblies.

This picture, however, is different for the SOM-PV network. As it turns out, PV cells which are more strongly connected to E cells in E_1_ (have a high value of *PV* → *E*_1_), connect weakly to neurons in E_2_, as the right side of the matrix is almost the inverted version of the left side of this matrix (Figure 8B). This fact indicates that the PV population has split into two sub-populations, each of which have more preference to connect strongly to one of the excitatory cell assemblies. We call the first half of the PV cells in the connectivity matrix-with the higher value of *PV* → *E*_1_-PV_1_ population. The second half with higher indices of PV cells-which in this case connect more strongly to E_2_ cells-are called members of the PV_2_ population. For the 1pv network, there was no distinction between PV_1_ and PV_2_, but nevertheless, we used the same labeling for this network as well.

To understand the origin of the emergent tuned PV to E weights, we checked the correlation functions between PV_1_ average firing rate and E_1_ and E_2_ average firing rates respectively, and we super-imposed it on the symmetric STDP curve that we used in our network simulations. The cross-correlation function between PV_1_ and E_2_ has a big negative trough which makes the entire integral of the product of the correlation function and the STDP curve negative (Figure 8E). This results in the depression of the connection weights from PV_1_ to E_2_. However, the cross-correlation function between PV_1_ and E_1_ is mainly positive, and the interaction of this function with the STDP curve yields a positive quantity for the rate of change of the connections from PV_1_ to E_1_. This justifies the emergence of tuned weights in the SOM-PV network. These correlation functions for the 1pv network are almost identical 8D). Hence, there is no preference for the connections from PV onto E_1_ or E_2_.

The 2PV network already has two distinct PV populations. Therefore, we did not sort the indices of the PV cells in this case. The PV to E connectivity matrix has a clear block structure, in which the weights from PV_1_ onto E_1_ and the weights from PV_2_ onto E_2_ are bigger than the other counterpart weights (Figure 8C). This tuned connection from PV_*x*_ to E_*x*_ can similarly be explained by examining the correlation functions between PV and E activity. The correlation function between PV_1_ and E_1_ is mostly positive, and this results in the potentiation of the inhibitory synapse connecting these two populations. However, the correlation function between PV_1_ and E_2_ is mainly negative, and this explains weaker connections from PV_1_ to E_2_.

So far, we have examined the correlation structure between the PV sub-populations and the excitatory cell assemblies. To shed light on the reason behind the difference in correlation functions between PV populations and the E populations in the SOM-PV network, we dissected the projection weights from the SOM populations onto the PV sub-populations. It turned out that SOM_1_ projected weakly onto PV_1_, but more strongly onto PV_2_ (Figure 8G). The same pattern of connectivity was observed for the total projection weights from SOM_2_ onto PV_2_ and PV_1_, respectively. This observations explains that when SOM_1_ neurons are active, the strong projection weights from this population onto E_2_ and PV_2_ suppresses both populations simultaneously. When SOM_1_ is relatively silent, both E_2_ and PV_2_ will get the chance to be activated together. As a result, PV_2_ and E_2_ will be correlated. The same holds true for the correlation structure between PV_1_ and E_1_. Since E_1_ and E_2_ are anti-correlated, PV_1_ and E_2_ (also PV_2_ and E_1_) will also represent negative correlation. This explains the correlation structure in Figure 8F.

Since the initial bias from E_1_ to PV_1_, and from E_2_ to PV_2_ was the origin of the emergence of reciprocal inhibitory connections in the 2PV network, we checked those random connections in the SOM-PV network as well. There was a slight bias in the overall excitatory connections from E_1_ to PV_1_ (Figure 8H), however, the distributions of the overall excitatory weights onto each subgroup of PV were not as distinguishable as the distributions of the inhibitory projection weights from SOM onto the PV subpopulations (Figure 8G). Hence, for the SOM-PV network, the main cause of the emergence of tuned PV to E connections were the heterogeneity in random connections from SOM to PV populations.

The distribution of the projection weights from SOM_1_ onto PV_1_ is not identical to the distribution of projection weights from SOM_1_ onto PV_2_ because of the random Bernoulli distribution of connections (Figure 8G). This fact can by chance break the symmetry in the total projection weights between these two distinct inhibitory subtype. This slight break in symmetry associates the PV neurons to either of the E assemblies. For the SOM-PV network considered here, slight symmetry breaking in the projections from E to PV does not cause a big difference in the distribution of the weights, and therefore, should not play as important role as the SOM to PV projections. In our mean field analysis, we assumed a slight bias in the probability of connections from SOM_1_ to PV_2_, and a bias from SOM_2_ to PV_1_, represented as *E* in equation (30). As derived in the methods section (equation (38)), the difference between the weights from PV_1_ to E_1_ and the weights from PV_1_ to E_2_ scales linearly and positively with *w*_11_ - *w*_12_ (difference between the weights within an excitatory assembly and between the two excitatory assemblies). Our network simulations result confirms this prediction, as illustrated in Figure 8I, where the difference between the tuned and untuned PV to E weights are plotted against the difference between the weights within and between the excitatory assemblies. This plot suggests an almost linear function approximation for the relation between these two variables. This implies that as long as assemblies are being learned (*w*_11_ - *w*_12_ > 0), there should emerge a tuned connection from the PV sub-population-that receives slightly stronger projection from the antagonistic SOM population-onto its corresponding excitatory assembly which also receives a strong projection weight from the same SOM population.

## Discussion

Cell assemblies are thought to be the building blocks of neuronal dynamics which map neuronal activities to behaviour [10, 7]. These coactivated groups of neurons are responsive both during spontaneous and stimulus conditions [6], and are believed to appear as a result of neuronal correlations under Hebbian plasticity [10] (also see [52]). While there has been some theoretical work to understand the role of inhibition in the formation of assemblies [13, 12, 11], there is an emerging experimental interest in understanding the contribution of inhibition assembly formation [26, 53, 54]. It is particularly interesting to study the role of inhibition in correlative learning because it is known that inhibition decorrelates the activity of excitatory neurons in the balanced condition [23, 24]. However, the structure of correlation when different inhibitory subtypes are involved is not clear. Since inhibitory plasticity plays a role in the formation of assemblies, and since inhibition plays a crucial role in shaping network correlations, it is important to understand how different inhibitory subtypes shape their connections to facilitate learning.

In [55], the authors reported a symmetric STDP function for the connections from SOM to pyramidal neurons, and a depressive STDP curve for the connections from PV to CA1 pyramidal neurons in the Hippocampus. Our experimental findings demonstrate that in the OFC, the STDP curve for the connections from PV to E cells follows a symmetric rule with potentiation of the synapse for small time lags between pre and post synaptic spikes. It is easy to show that this kind of STDP curve can result in a homeostasis rule to regulate the firing rate of the post-synaptic neuron, conditioned that upon the firing of the inhibitory neuron, the synapse depresses. Regulating the postsynaptic firing rate brings stability to the system, and prevents runaway dynamics, particularly during training.

We showed, for the first time, that the STDP curve for the connections from SOM to E cells in the OFC follow an asymmetric Hebbian learning rule. In our large scale network simulations, we demonstrated that if the excitatory neurons have a tuned connection to SOM cells, then the causality in the STDP rule can weaken the synapse from SOM to its driving excitatory population, and potentiate the synapse from SOM to the competitor excitatory population. This results in a di-synaptic pathway with lateral inhibition between the excitatory assemblies. Lateral inhibition results in more pronounced competition between assemblies [56, 57]. We demonstrated both in network simulations, and in our theoretical framework, that the existence of lateral inhibition can amplify the correlation functions between excitatory neurons.

Whether excitatory neurons have a tuned connectivity profile to inhibitory neurons in the OFC is not clear. In other cortices such as the visual cortex, there are some evidence for the existence of tuned excitatory to PV connections[58], and tuned excitatory to SOM connections [59, 60]. Also, whether inhibitory subtypes in the OFC are tuned to a specific input feature is not known. In the visual cortex of mice, it has been claimed that despite the fact that excitatory neurons are sharply selective for certain input features, Gabaergic interneurons are less selective and broadly tuned [61, 62], and they do not form subnetworks of coacivated groups [63]. However, other studies report that PV cells in the visual cortex are integrated into subnetworks of excitatory cells with similar visual selectivity [58, 64]. Considering this ambiguity, we considered 3 different networks: the 1pv network with a shared pool of PV neurons, without any tuned connections, the 2PV network with two distinct groups of PV cells that received stronger projection weights from their feature-specific excitatory population, and the SOM-PV network with tuned connections from the excitatory populations to their specific SOM population. The SOM-PV network also had a PV population which received no tuned input from the excitatory subnetworks. In our network simulations, we observed that during and after training, the correlation structure for neurons in the same assembly was the strongest in the SOM-PV network, and hence assemblies were formed faster. Correlations between excitatory neurons within each group were weak in the 2PV network. The correlation structure was reversed between the excitatory neurons that belonged to different assemblies. Increased correlations can come about not only through lateral inhibition, but also through reciprocal inhibitory to excitatory populations. The heterogeneity in random connections from SOM to PV cells provided a substrate for correlations between a group of PV and individual excitatory assemblies such that neurons which received strong inhibition from SOM were more correlated. This resulted in tuned PV to cell assembly connections due to the symmetric shape of the STDP curve for PV to E connections, and the fact that these connections should maintain a constant firing rate. These reciprocal connections also play an important role in shaping correlations between excitatory assemblies. The 2PV network with biased connections from E to PV cells also developed reciprocal connections from PV back to the driving excitatory assemblies. However, due to lack of strong competition and weaker correlations, assemblies did not form as quickly as those in the SOM-PV network. Existence of tuned PV to E connections and subnetworks of excitatory and PV cells with relatively strong connections were reported earlier in the visual cortex [65, 64, 58], and our network simulations and theoretical analysis confirms this finding. Interestingly, in a network with homeostasis but no tuned excitatory to inhibitory connections, like the 1pv network we studied here, no tuned PV to E connections emerge. The reason is that the correlation structure between PV and excitatory populations is not different across PV neurons.

In our study, the amplified cross-correlation function between excitatory cell assemblies implies promoted competition, which results from SOM plasticity onto excitatory neurons. While the homeostasis provided by PV plasticity can also impose competition between assemblies [13, 12, 11], we showed that the strength of competition is enhanced when SOM plasticity contributes to learning. However, the amplitude of fluctuation can increase significantly, and beyond control, if there is no homeostatic mechanism to maintain a fixed firing rate for the excitatory neurons. We conclude that PV plasticity is responsible for providing circuit stability, while SOM plasticity promotes competition. This division of labor is necessary because one source of inhibition alone cannot be responsible for both functions in an adequate way.

We showed throughout the paper that increased correlation between excitatory neurons due to SOM plasticity contribution can result in faster formation of assemblies. Increased speed of learning is critical for animal survival, and this improved cognitive function results from SOM inhibitory subtype plasticity. Also, considering different network structures, we showed that after the training signal was removed, in the presence of SOM inhibitory subtype, assemblies could maintain their structure for a longer time. More precisely, for a given short training period, the network without SOM inhibitory subtype lost its assembly structure completely, however, the network including this subtype recovered the assembly structure shortly after training. This indicates that SOM plasticity helps assemblies to be more robust to perturbations after training, and protects them against decorrelative factors, such as global input projections, or spontaneous brain dynamics.

In summary, based on our experimental results for the spike timing dependent plasticity curves for the connections from different inhibitory subtypes-mainly SOM and PV neurons-onto excitatory cells, we conclude that while PV cells provide circuit stability, SOM cells are the main contributors to provide competition between excitatory cells, which in turn accelerate assembly formation and protect assemblies from degeneration. One way to test our predictions would be to block the activity of SOM cells and compare the correlation structure between excitatory neurons before and after this blockage. According to our study, if SOM activity is removed from the network dynamics, and if the firing rate of PV does not change much as a result of this dysfunction, then the correlation between the excitatory neurons should decrease. Another prediction is that animals’ behavior in odor discrimination tasks should be impaired if the activity of SOM cells are blocked (either during or after training the animal with new odors) because the structure of assemblies fade away very quickly when SOM activity is blocked.

## Methods

### Experimental details

#### Mice

SST-Cre (B6:Sst<tm2.1(cre)Zjh>/J) and PV-Cre (B6.129P2-Pvalbtm1(cre)Arbr/J) mice [66] that express cre-recombinase were crossed with Ai32 mice (B6;129S-Gt ROSA)26Sortm32 (CAG-COP4*H134R/EYFP)Hze/J) [67] to express channelrhodopsin (ChR2) in somatostatin or parvalbumin interneurons, respectively. All mice were from Jackson Laboratory. Slice preparation: Coronal brain slices of orbitofrontal cortex (OFC) located 2.845-2.245 mm anterior to Bregma (Allen Brain Atlas) were prepared from mice of both sexes that were P20-28 days old. The mice were anesthetized with isoflurane and decapitated. The brain was removed from the skull and immersed in ice-cold oxygenated (95% O2-5% CO2) ACSF solution (in mM: 125 NaCl, 2.5 KCl, 25 NaHCO_3_, 1.25 NaH_2_PO_4_, 1.0 MgCl_2_, 25 Dextrose, 2.5 CaCl_2_) (Sigma-Aldrich, USA). Coronal slices were 300 m thick and cut using a vibratome (Leica Biosystems) in ice cold ACSF. The slices were transferred to warm ACSF (36 C) for 30 min and then rested at 20-22 C for 45-60 min prior to recording in the same conditions. All procedures used in this study were approved by The University of Pittsburgh IACUC.

#### Electrophysiology

Recordings were obtained from L2/3 excitatory neurons with input resistance <250 MΩ visualized using infrared differential interference contrast microscopy (IR-DIC, Olympus). For all recorded neurons, intrinsic subthreshold properties such as input resistance, and time constant were obtained using a series of hyperpolarizing and depolarizing current steps (10 pA, 1 s duration). Whole cell, voltage and current clamp recordings were performed using a MultiClamp 700B amplifier (Molecular Devices, Union City, CA). Data were low pass filtered (4 kHz) and digitized at 10 kHz using an ITC-18 (Instrutech) controlled by custom software (Recording Artist, *https://bitbucket.org/rgerkin/recording-artist*) written in IgorPro (Wavemetrics). Recording pipettes (5-10 MΩ were pulled from borosilicate glass (1.5 mm, outer diameter) on a Flaming/Brown micropipette puller (Sutter Instruments). The series resistance (<20 MΩ) was not corrected. The intracellular solution consisted of (in mM) 130 K-gluconate, 5 KCl, 2 MgCl_2_, 4 ATP-Mg, 0.3 GTP, 10 HEPES, and 10 phosphocreatine, 0.05% biocytin. Since post-synaptic spike responses are required for STDP induction, we could not use Cesium or QX-314 in the recording pipette. The reversal potential for inhibition was found for each neuron to be between -75 to -70 mV. All voltage clamp recordings were performed with a holding potential between -50 to -60 mV to maximize inhibitory currents but maintain a subthreshold membrane potential in the PN. To isolate GABA_*A*_ receptor mediated inhibition, the GABA_*B*_ receptor antagonist, Saclofen (20-40 M), was included in the bath.

#### STDP protocol

The STDP induction protocol was applied with 10 min of patching the PN. Baseline IPSCs were light-evoked from either PV or SOM interneurons every 30 s and recorded in voltage clamp for 5 min prior to STDP induction. Recordings were then switched to current clamp to evoke action potentials in the post synaptic PN during the STDP protocol. To induce STDP, a single action potential (AP) was evoked by current injection at the soma (50 *µ*s pulse width). The AP was paired with light-evoked IPSP at a consistent time interval with respect to the AP. We applied this pairing protocol every 4 sec (0.25 Hz) for 4 min. The interval is defined as the difference in time (ms) between the AP and the light evoked IPSP. Following the STDP protocol, IPSCs were again evoked every 30 s for up to 1 hour. In all cases, series resistance and input resistance were monitored throughout baseline, induction and post-induction testing. Neurons with deviations greater than 25% from baseline were excluded from analysis.

#### Optogenetic Stimulation

Shutter controlled full field light stimulation of blue light (473 nm) provided by a mercury lamp was delivered through the epifluorescence pathway of the microscope (Olympus) using a water-immersion objective (40x). The duration of the light pulses was 5 ms, light intensity (3-4 mW) was adjusted to induce reliable, single spike responses in neurons expressing ChR2 (Figure 2B1,C1).

#### Data analysis

Inhibitory strength was taken as the peak amplitude of the IPSC within 10 ms of IPSC onset. Since these are population IPSCs, the rising slope of the IPSC and the integral of the IPSC were also analyzed. The results did not differ from the main findings reported in this study (data not shown). To assess the time course of plasticity, IPSC amplitudes pre and post induction were normalized to the average baseline amplitude. To assess the change in synaptic strength following the STDP protocol, to took the average normalized IPSC amplitude between 25-30 min post induction (10 IPSCs). We chose this time point to minimize any influence of recording drift across neurons. However, in the majority of neurons the level of synaptic plasticity measured at 30 min was maintained until the end of recording (1 hr).

#### Statistics

All data is presented as mean SE. Statistical tests were performed using two tailed, one or two-sample, unpaired Students t-test as appropriate. In cases of small sample sizes the non-parametric tests Mann-Whitney U-test (MWU) test was used.

### Network simulations

In all networks we studied (SOM-PV, 1pv, and 2PV networks), 80% of the neurons were excitatory and the remaining 20 % were inhibitory. We used Leaky Integrate-and-Fire (LIF) neuron models with delta shape PSC. The membrane potential dynamics for the LIF neuron was a low pass filter of first order with the dynamics

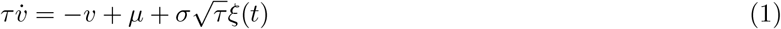

where *µ* and *σ* represent the mean and the standard deviation of the fluctuations imposed on the neuron as a result of external input. The variable *ξ*(*t*) is a normal Gaussian white noise.

Each excitatory population was composed of *N*_*e*_ neurons, and each Somatostatin population had *N*_*s*_ neurons. The inhibitory PV population was composed of *N*_*i*_ neurons for the SOM-PV network, and it had 2.5 times more inhibitory neurons for both 1pv and 2PV networks. The probability of connection between excitatory neurons was *p*, and every other probability of connection was 4*p*. As such, every two neurons were connected according to a Bernoulli distribution with parameters *p* or 4*p*.

The non-plastic excitatory weight to PV neurons is represented by *J*, which in our study took the value of 0.06 mV, and the fixed inhibitory weights are −*gJ*.

For the SOM-PV network, we chose tuned excitatory to SOM connections by strengthening the weights by a factor of 2, meaning that projections from the excitatory populations to their corresponding SOM population was 2*J*. Also, the probability of connection from each excitatory assembly to their corresponding SOM population was *p*_*t*_ = 2(4*p*)*/*3 = 8*p/*3, in which *p*_*t*_ represents tuned probability of connections. The tuned excitatory to PV connections in the 2PV network also followed the same method.

Each neuron in the excitatory and PV population received spikes from an independent Poisson process with a firing rate of *σ*^2^ when there was no training. During training however, neurons received spikes from independent external sources with a firing rate of 0.7*σ*^2^, and a shared source of spikes with a rate equal to 0.3*σ*^2^.

The network simulation time resolution and also the synaptic transmission delay between neurons were chosen to be 0.1*ms*.

### Plasticity equations

The STDP curve for the excitatory connections was

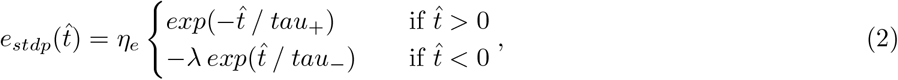

The STDP curve for the connections from SOM to excitatory neurons was

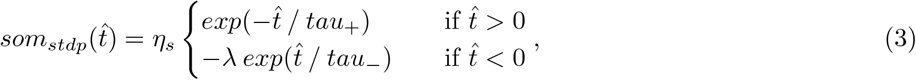

where we chose *τ*_+_ = 30 ms and *τ*_−_ = 30 ms for SOM stdp, and *τ*_+_ = 15 ms and *τ*_−_ = 30 ms for the excitatory stdp. The factor *λ* = 0.53.

The symmetric STDP curve for the connections from PV to E was

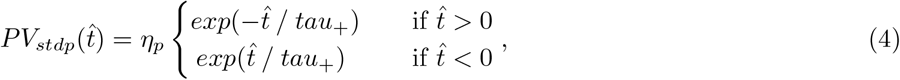

The time constant of the STDP curve for the PV to E connections was *τ*_+_ = 20 ms. The time constant of learning for the excitatory to excitatory connections were *η*_*e*_, and the equivalent quantity for the connections from SOM to E were *η*_*s*_.

### Mean-field equations

The dynamic mean-field equations for individual neurons for the SOM–PV network are

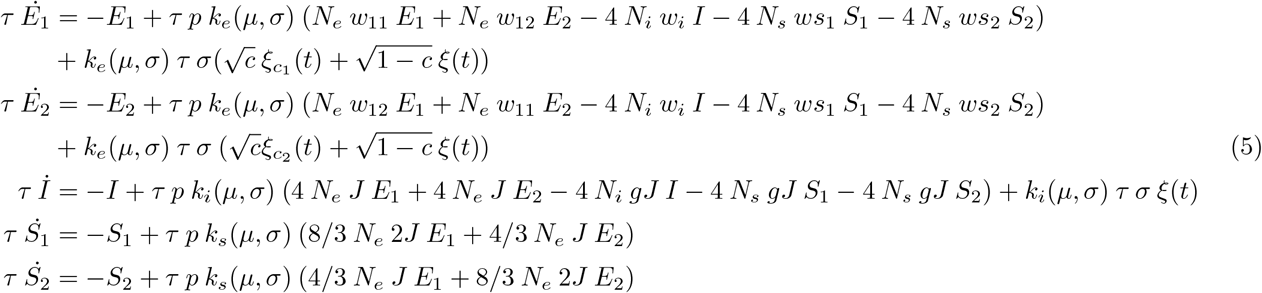

for the details of the derivation see the supporting information.

## Supporting Information

### Coupled rate and weight dynamics

Here, we study the interactions between dynamics of leaky integrate-and-fire (LIF) neurons and the evolution of weights between neurons in different assemblies, as well as the inhibitory weights to the excitatory populations. The membrane potential dynamics for an LIF neuron is a low pass filter of first order with the dynamics

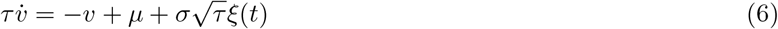

where *ξ*(*t*) is a normal Gaussian white noise.

In order to get neuronal correlation functions, it is required to linearize the dynamics of individual neurons around their operating stationary firing rates. A transfer function that relates the stationary input to the firing rate of the neuron is obtained from the stationary solution of the Fokker-Planck equation to solve a first passage time problem. The stationary firing rate r is obtained from *r* = *f* (*µ, σ*) where the transfer function *f* (.) follows

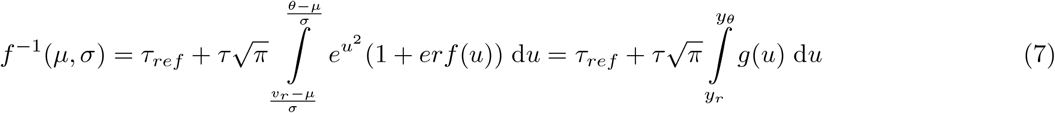

To linearize the transfer function around the operating point *r*\*, we take the derivative of both sides with respect to *µ*. This results in

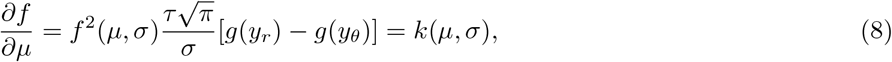

where *k*(*µ, σ*) represents the slope of the *f* − *I* curve at the linearization point, and 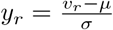 and 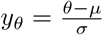.

We choose *E*_1_ and *E*_2_ to represent the dynamic firing rates of individual neurons in the excitatory assemblies, *I* to represent the fluctuating activity of one neuron in the PV inhibitory population, and *S*_1_ and *S*_2_ to stand for the dynamic firing rates of the somatostatin neurons. The operating point for excitatory, PV and SOM neurons are 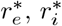, and 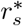, respectively (the stationary solution comes from solving coupled Fokker-Planck equations). The connection weights from population j to i are *w*_*ij*_. The variable *w*_*i*_ indicates the weight from PV to excitatory neurons. The variable *ws*_*i*_ represents the connection weight from a SOM neuron to an excitatory neuron in assembly *i*.

The dynamic mean-field equations for individual neurons are

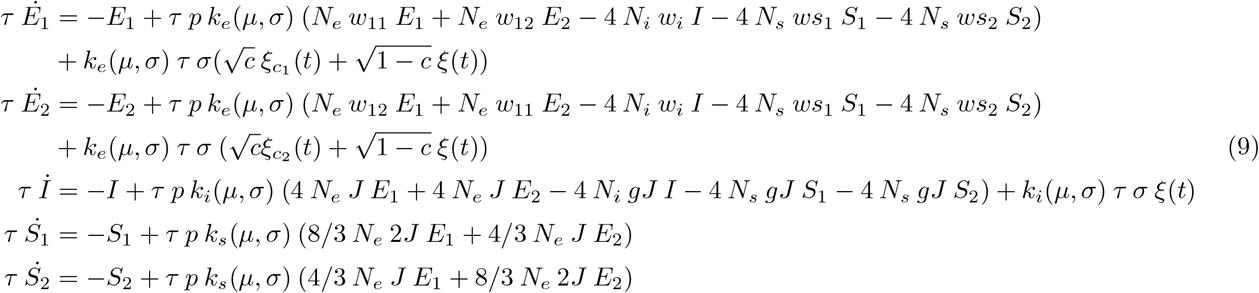

which in matrix form can be represented as

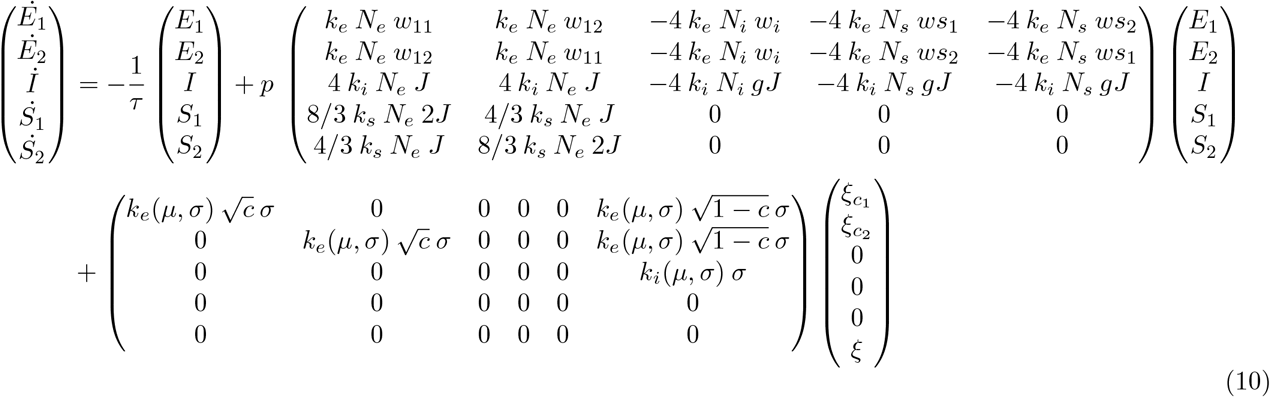

The fluctuation of the rate dynamics around the fixed point can be written in the following general form:

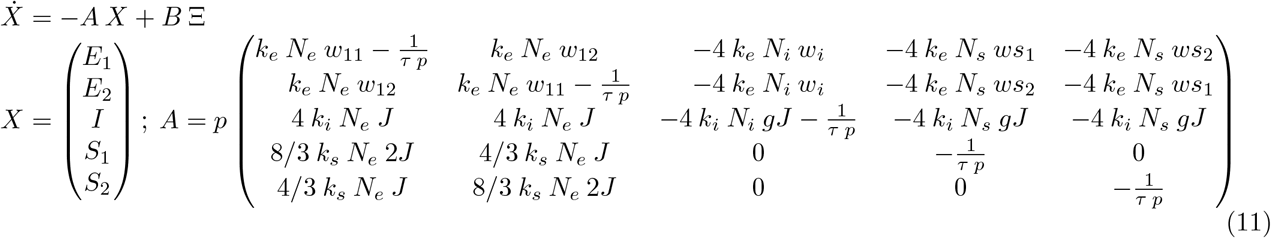

The auto-covariance matrix for *X* is

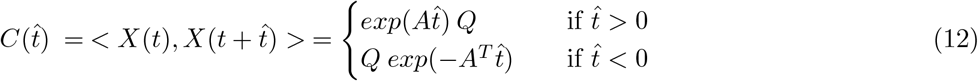

and *Q* is the solution to the following Lyapunov equation

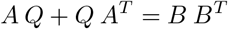

The dynamics of the weight evolution is slow. In order to obtain a governing equation for individual weight dynamics, we need to calculate the integral of the product between the stdp curve of the synapse, and the cross correlation between the pre and post synaptic firing rates.

From Kempter et.al, the dynamics of the weights follow

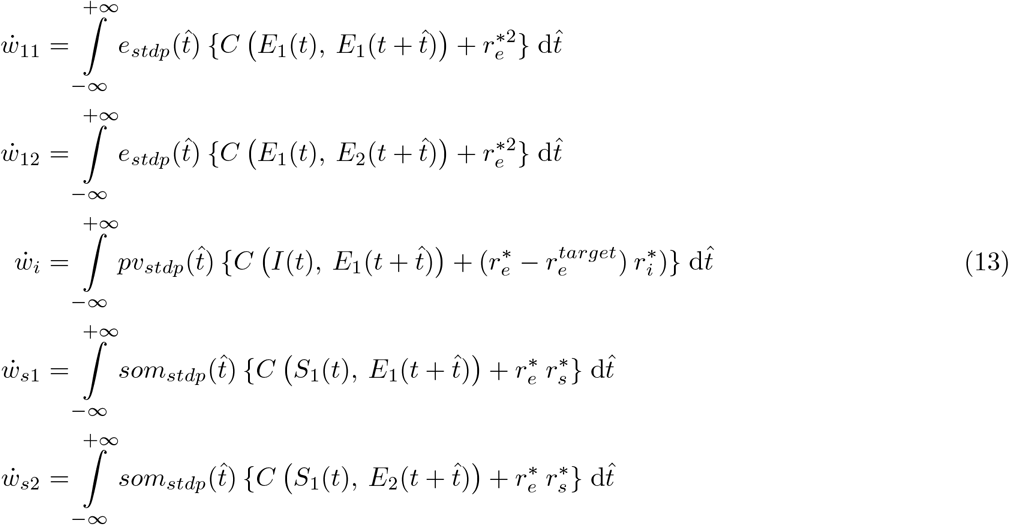

We assume that the operating point does not change when the weights are changing, i.e *r*\* = *constant*. The challenge is to write the weight dynamics (13) in a self-consistent way, such that it only depends on the weights of the network. The correlation function (12) can be calculated semi-analytically (not analytically due to the existence of *Q*), using the Laplace transform. Since the impulse response of the general system 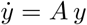 is *exp*(*A t*) in time domain, and (*s***I** - *A*)^−1^ in the Laplace domain (*s* is the complex variable defined in the Laplace transform, and **I** is the identity matrix), the matrix exponentials in equation (12) can easily be replaced by (*s***I** - *A*)^−1^ and (*s***I** - *A*^*T*^)^−1^, respectively, when needed. This can also help to evaluate the integrals in equation (13) due to the fact that the product of a linear system and an exponential function causes a shift in the Laplace domain. Recruiting this transform, equations (12) and (13) can be rewritten as

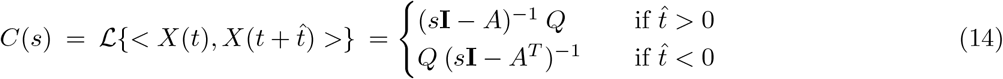

and the element on row *i* and column *j* of matrix *C*(*s*) will be represented by *C*_*i,j*_(*s*). Now, equation series (13) can be written as the following form, where the variable *s* will be replaced by 0 in order to get the values of the integrals in (13):

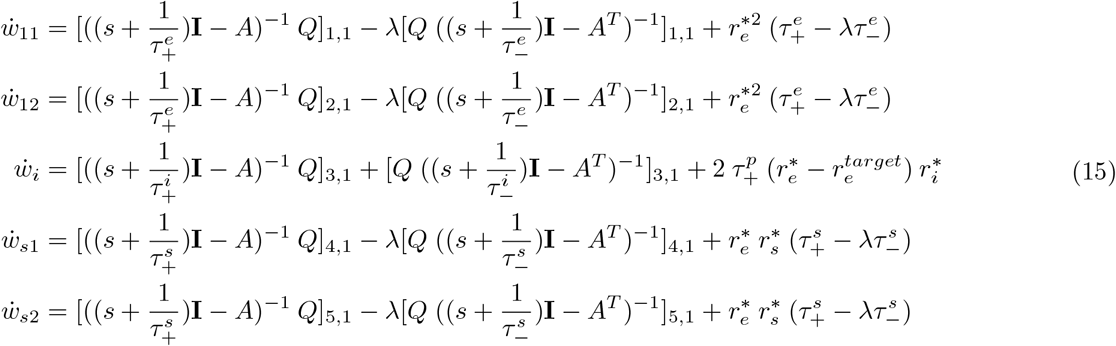

In above, 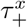 and 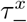 represent the exponent for the positive and negative domains of the stdp curves for the neuron type *x ϵ* {*e, i, s*}.

### Dimensionality reduction for the SOM-PV network

In order to reduce the dimensionality of the system such that a phase portrait plot in *w*_11_ - *w*12 plane is possible, we rewrite the dynamics of the rates in the Laplace domain:

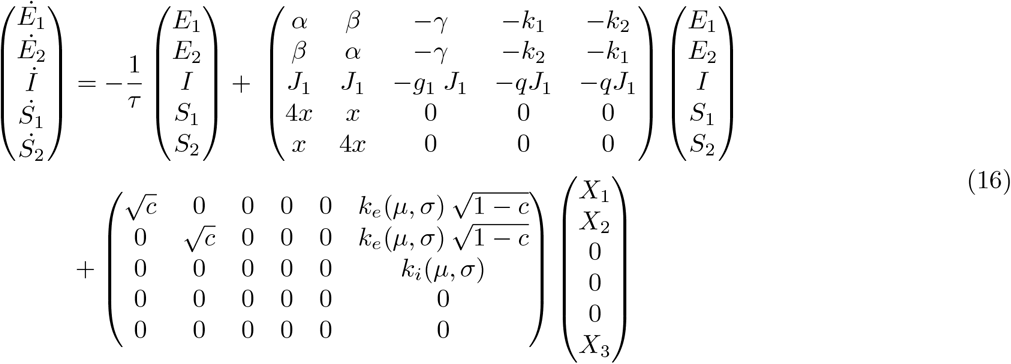

where 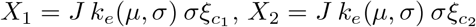, and *X*_3_ = *J σξ*.

*α* = *k*_*e*_(*µ, σ*) *p N*_*e*_ *w*_11_,

*β* = *k*_*e*_(*µ, σ*) *p N*_*e*_ *w*_12_,

*γ* = −4 *p k*_*e*_ *N*_*i*_ *w*_*i*_,

*k*_1_ = −4 *p k*_*e*_ *N*_*s*_ *ws*_1_,

*k*_2_ = −4 *p k*_*e*_ *N*_*s*_ *ws*_2_,

*J*_1_ = 4 *p k*_*i*_ *N*_*e*_ *J*,

*g*_1_ = *N*_*i*_*/N*_*e*_ *g*,

*q* = *N*_*s*_*/N*_*e*_ *g*,

*x* = (0.8*/*3) *J N*_*e*_ *k*_*s*_(*µ, σ*).

In the Laplace domain, the equations can be written in the following 3 dimensional form:

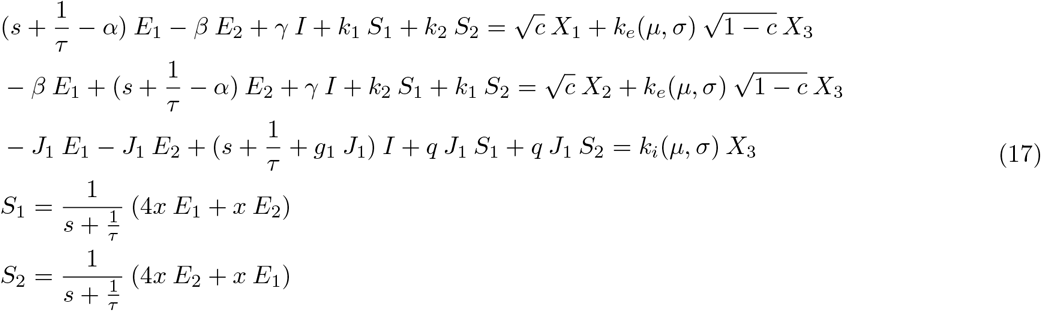

which yields

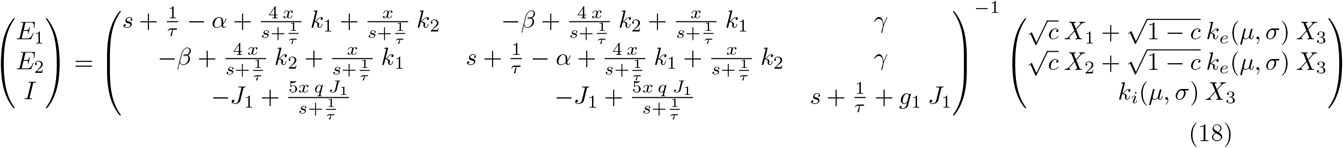

The inverse of the matrix in equation (18) can be represented as

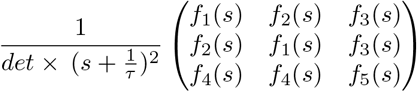

where

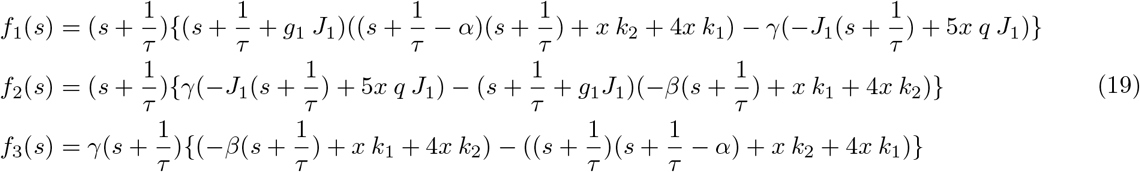

Therefore, the solutions for *E*_1_(*s*) and *E*_2_(*s*) are

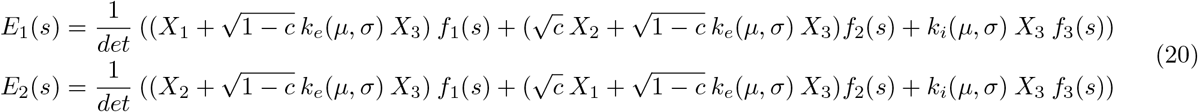

where *det* is the determinant of the characteristic matrix in equation (18) (or the product of (*s* − *eigenvalues*) of the matrix A in the 5 dimensional system)

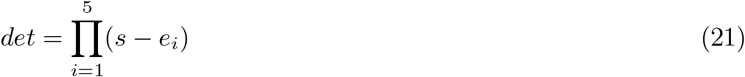

The correlation functions in the Laplace domain are

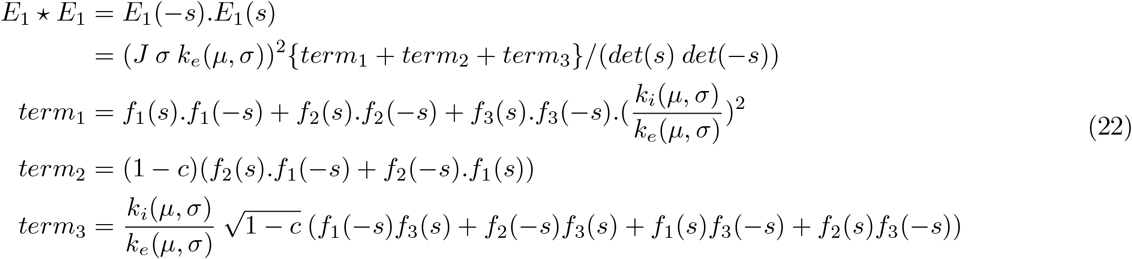

and

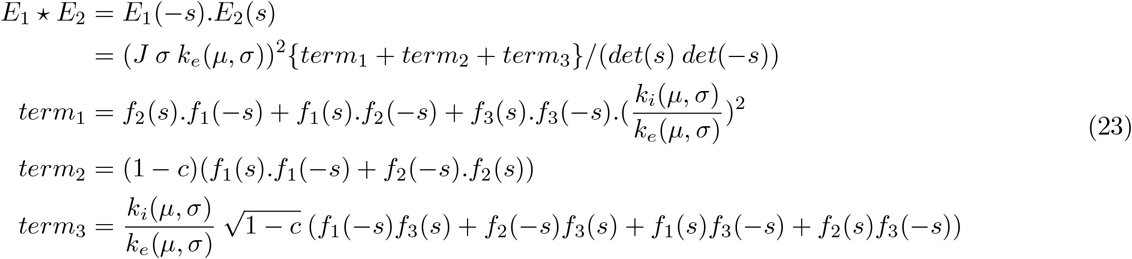

However, these functions include the correlation functions for both *t* > 0 and *t* < 0. Due to the symmetry of these functions around *t* = 0, we calculate the functions for *t* > 0. To do so, we need to use the technique of partial fractions and keep the terms with components of *det*(*s*) only, and neglect those with components of *det*(-*s*). This results in

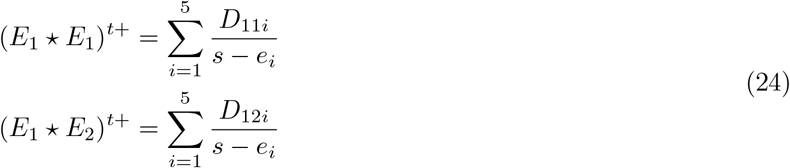

in which *D*_11*i*_ and *D*_12*i*_ are obtained by replacing *s* in *E*_1_ *E*_1_ and *E*_1_ *E*_2_ by *e*_*i*_, respectively. According to equation (13), the dynamics of the weights in the Laplace domain are

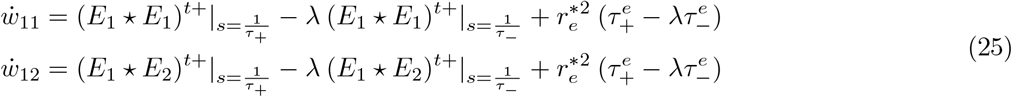

In the equations above, the term *γ* contains the plastic inhibitory weight from PV to E neurons. Because homeostasis on the firing rate of E cells guarantees a constant rate, we can assume that the mean membrane potential of the E neurons are constant, and we call this term *const*. From the dynamics of the LIF neuron (6), it is straightforward to replace *γ* by a function of other variables in the network:

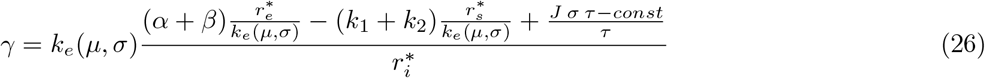

Moreover, the dynamics of the inhibitory weights from SOM to E neurons can be written as a function of *w*_11_ and *w*_12_ (or in other words, *α* and *β*). To get this relation, we use both equations (17) and (15). It is clear that

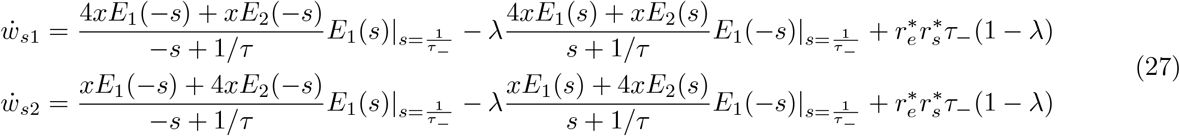

Since in the equation above the terms *E*_1_(−*s*)*E*_1_(*s*) and *E*_2_(−*s*)*E*_1_(*s*) appear, it is conceivable to assume that *w*_*s*1_ and *w*_*s*2_ are linear functions of *w*_11_ and *w*_12_.

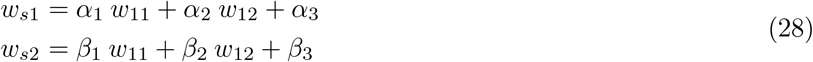

Following this idea, we replace *k*_1_ and *k*_2_ with a linear combination of *α* and *β*:

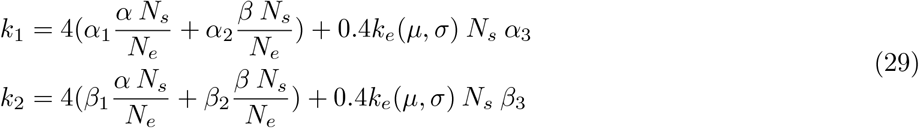

Using equations (29) and (26) in equation (25), we obtain a system that can describe the weights within and between assemblies entirely as a function of *w*_11_ and *w*_12_, and other constants of the system. The same approach can be used to reduce the dimensionality in the 1pv network. In that case, there is no *k*_1_ and *k*_2_, and only *γ* has to be replaced by *α* and *β*.

### Tuned PV to E connections

In order to study the emergence of tuned inhibitory weights from *PV* to *E*_1_ and *E*_2_, we assume that in equation (16), *S*_1_ projects slightly more strongly to *E*_2_ than to *E*_1_. To model this symmetry breaking, we assume that *E* represents the amount of asymmetry in projections from *SOM* to *PV* populations. The PV population, therefore, needs to be split into two different sub-populations, called *I*_1_ and *I*_2_. Since in this case we have a 6 dimensional system, we represent the new variables by a small symbol on top of the old variables. The new firing rate dynamics in the time domain would be

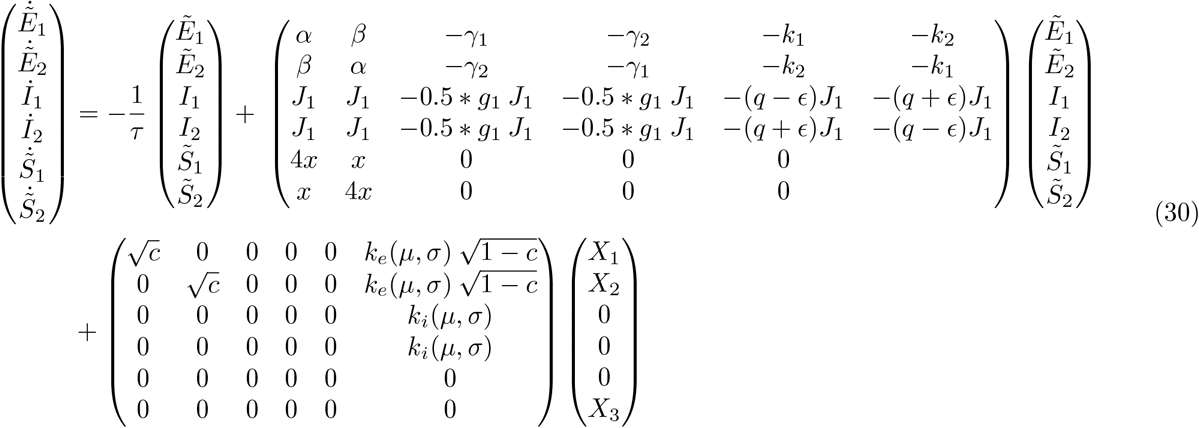

Note that the factor 0.5 on the third column and the third row (and similarly other 0.5 factors) are due to the fact that in these new equations, *I*_1_ and *I*_2_ have half of the population size for *I* in equation (16). More precisely, the dynamics of *I*_1_ and *I*_2_ are

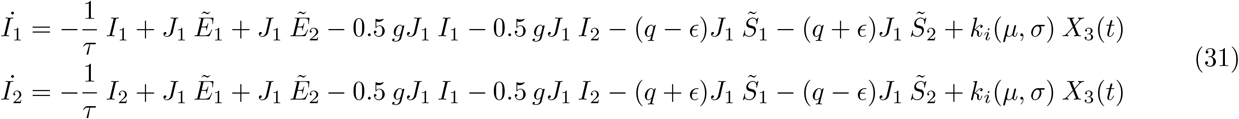

Summing up and subtracting the equations in (31), one obtains

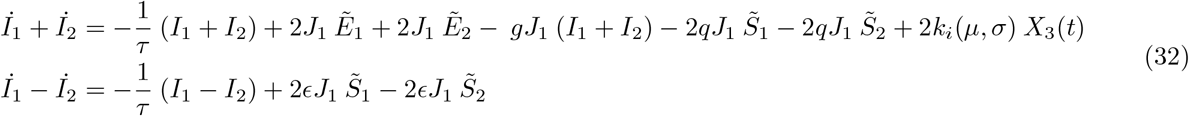

Comparing the first equation in (32) with the third equation in (16), one concludes *I*_1_ + *I*_2_ = 2*Ĩ*. Here, *Ĩ* is a variable similar, but not equal to *I* in (16). Since this variable does not play role in the final result, we are not interested in getting its precise temporal dynamics, and we will leave it as an abstract variable.

The new equations for the firing rates of the excitatory neurons are

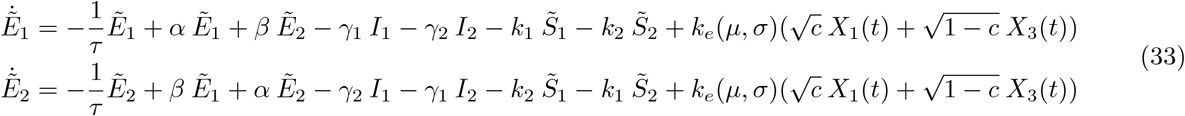

The last two equations in (17) hold in this new system. Usin the formula which relate 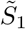 and 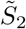 to 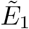 and 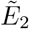, and also including our previous conclusion, we will get the following set of relations between *I*_*1*_, *I*_*2*_, *Ĩ*, 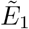 and 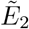 in the Laplace domain

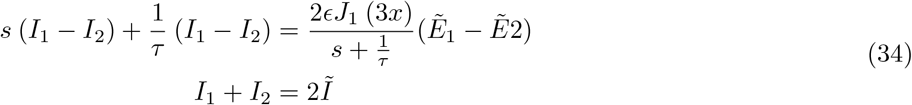

From the equation above, the solutions for *I*_1_ and *I*_2_ are

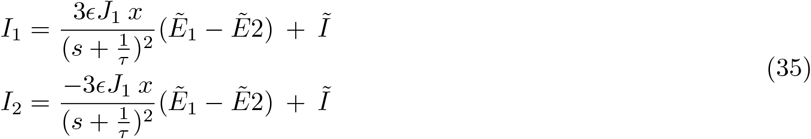

We are interested in the dynamics of *γ*_1_ and *γ*_2_, the inhibitory weights from *I*_1_ to *E*_1_, and *E*_2_, respectively. Using the formula in (13), the dynamics of *γ*_1_ and *γ*_2_ are

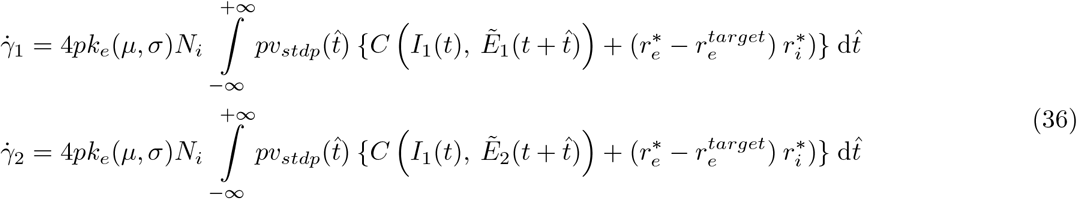

Since the second terms in the equation above are similar, we ignore their precise terms for the rest of our analysis, and will show them by *dc*. In the Laplace domain, the dynamic part of equation (36) are

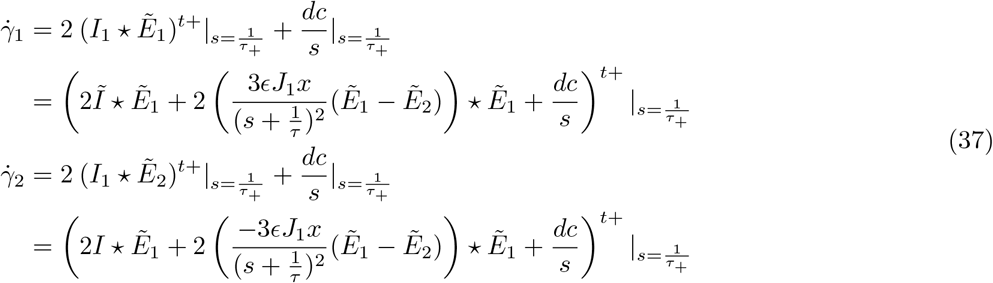

which results in

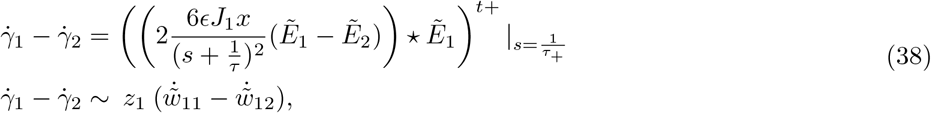

in which *z*_1_ is a positive scalar. Equation (38) indicates that as long as 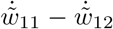 is positive, or in other words, as long as assemblies are being formed, 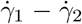 would be positive. This means that *γ*_1_ increases faster than *γ*_2_, and if the initial conditions for those variables were identical at the beginning of the simulation, after some time, *γ*_1_ > *γ*_2_. This shows the emergence of tuned *I*_1_ to 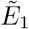 connections.

### Dimensionality reduction for the 1pv network

The dynamics of the 1pv network is simpler than the SOM-PV network. In this system, 3 dynamical variables *E*_1_, *E*_2_, and *I*, represent the dynamics of the firing rates for the two excitatory assemblies, and the shared inhibitory assembly. Moreover, there are 3 variables representing the within assembly weight dynamics (*w*_11_), between assembly weight dynamics (*w*_12_), and the plastic inhibitory weights from the PV neurons to the excitatory neurons (*w*_*i*_).

The dynamics of the firing rates can be written in the following form

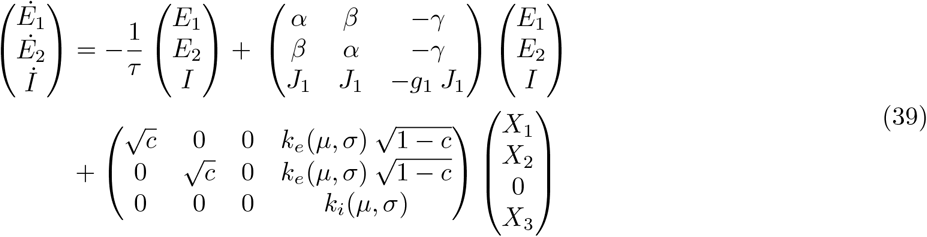

where 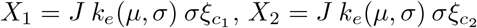, and *X*_3_ = *J σξ*.

*α* = *k*_*e*_(*µ, σ*) *p N*_*e*_ *w*_11_,

*β* = *k*_*e*_(*µ, σ*) *p N*_*e*_ *w*_12_,

*γ* = −4 *p k*_*e*_ *N*_*i*_ *w*_*i*_,

*J*_1_ = 4 *p k*_*i*_ *N*_*e*_ *J*,

*g*_1_ = *N*_*i*_*/N*_*e*_ *g*.

In the Laplace domain, the equations can be written in the following 3 dimensional form:

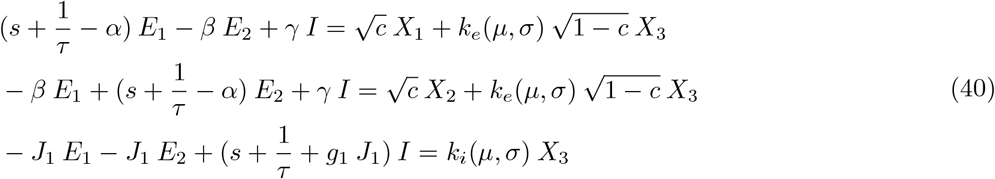

which yields

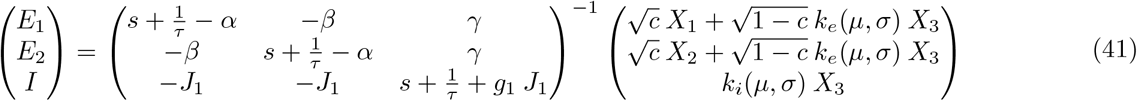

The characteristic equation of the system in equation (41) is:

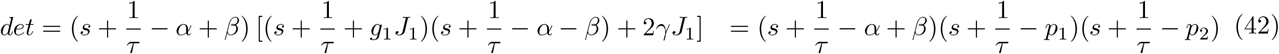

where *p*_1_ and *p*_2_ are 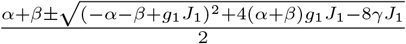

The inverse of the matrix in equation (41) can be represented as

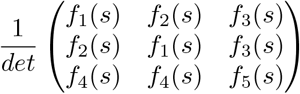

where

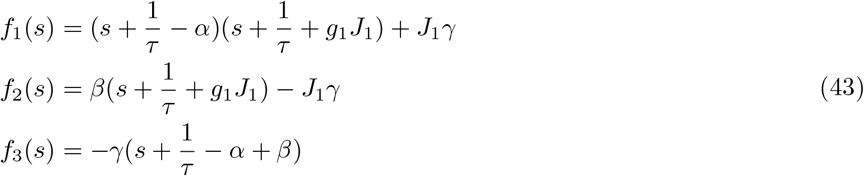

Therefore, the solutions for *E*_1_(*s*) and *E*_2_(*s*) are

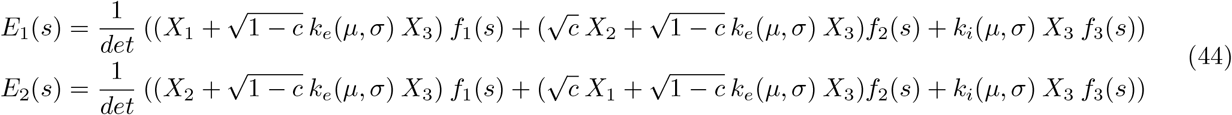

The correlation functions in the Laplace domain are

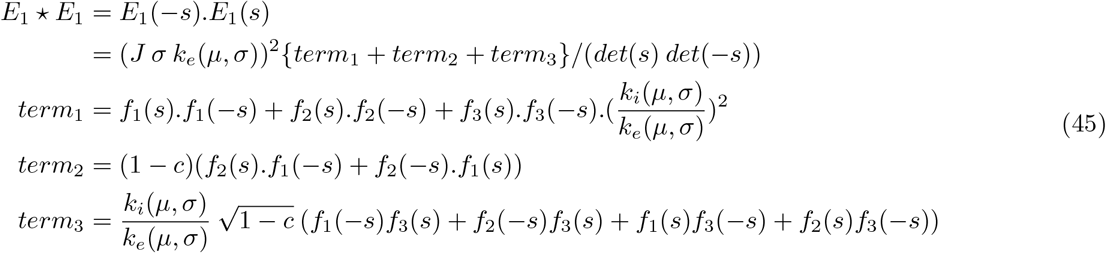

and

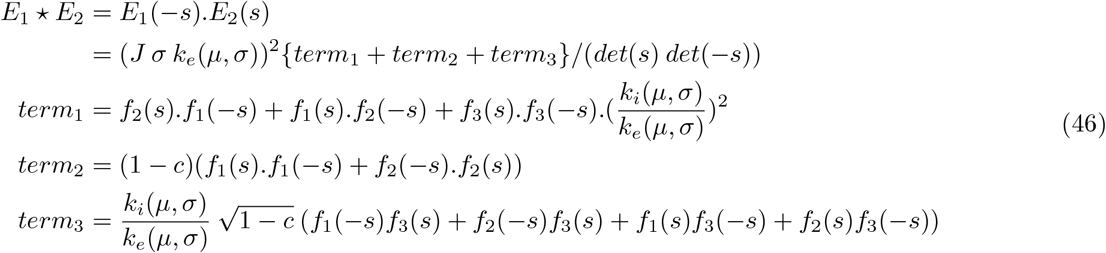

The solutions for *E*_1_ and *E*_2_, and hence the correlation functions are similar to those for the SOM-PV network; however, the functions *f*_*i*_(.), *i* = 1…5 are different in this system.

Similar to the solution for the SOM-PV network, the result in equation (46) includes the correlation functions for both *t* > 0 and *t* < 0. Due to the symmetry of these functions around *t* = 0, we calculate the functions for *t* > 0. To do so, we need to use the technique of partial fractions and keep the terms with components of *det*(*s*) only, and neglect those with components of *det*(−*s*). This results in

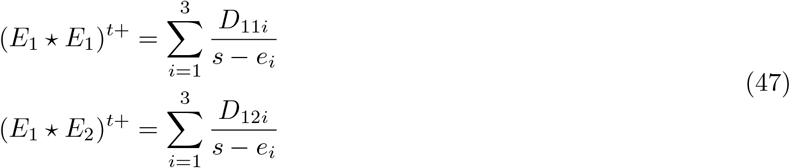

in which *D*_11*i*_ and *D*_12*i*_ are obtained by replacing *s* in *E*_1_ *E*_1_ and *E*_1_ *E*_2_ by the solutions of the characteristic equation (42), respectively. According to equation (13), the dynamics of the weights in the Laplace domain are

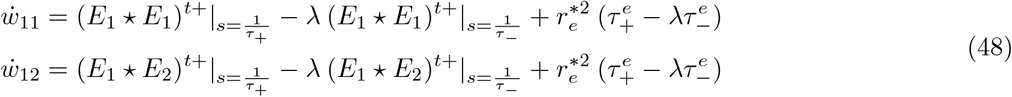

In the equations above, the term *γ* contains the plastic inhibitory weight from PV to E neurons. Because homeostasis on the firing rate of E cells guarantees a constant rate, we can assume that the mean membrane potential of the E neurons are constant, and we call this term *const*. From the dynamics of the LIF neuron (6), it is straightforward to replace *γ* by a function of other variables in the network:

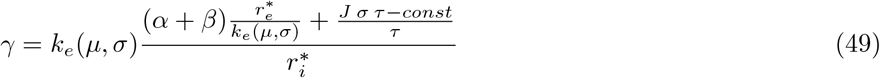

After this replacement, equation (48) describes the dynamics of *w*_11_ and *w*_12_ entirely based on these variables in a 2-dimensional system.

### Network parameters

Network parameters are represented in the following table.

| Variable | Value | Explanation |
| --- | --- | --- |
| $N_e$ | 400 | Number of excitatory neurons in each assembly |
| $N_i$ | 80 | Number of inhibitory neurons in the SOM-PV network |
| $N_i$ | 200 | Number of inhibitory neurons in the 1pv and 2PV network |
| $N_s$ | 60 | Number of inhibitory neurons in each SOM population |
| $J$ | 0.06 mV | Epsp amplitude for fixed connections |
| $g$ | 10 | Scalar for the fixed IPSP amplitudes |
| $I_e$ | 200 pA | External DC current to the SOM neurons |
| $\theta$ | 20 mV | Threshold for the membrane potential |
| $\tau$ | 20 ms | membrane potential time constant |
| $C$ | 250 pF | Capacitance of the membrane potential |
| $p$ | 0.1 | probability of connection between excitatory neurons |
| $p_t$ | 0.8/3 | probability of tuned E to SOM connections in the SOM-PV network |
| $p_t$ | 0.8/3 | probability of tuned E to PV connections in the 2PV network |
| $\tau_-$ | 30 ms | time constant of the depression side of SOM stdp |
| $\tau_-$ | 30 ms | time constant of the depression side of the excitatory stdp |
| $\tau_+$ | 30 ms | time constant of the potentiation side of the SOM stdp |
| $\tau_+$ | 15 ms | time constant of the potentiation side of the excitatory stdp |
| $\tau_+$ | 20 ms | time constant of the symmetric PV to E stdp |
| $\eta_e$ | $0.0025 \text{ ms}^{-1}$ | time constant of $E \rightarrow E$ connections |
| $\eta_s$ | $0.01 \text{ ms}^{-1}$ | time constant of $SOM \rightarrow E$ connections |
| $\eta_p$ | $0.01 \text{ ms}^{-1}$ | time constant of $PV \rightarrow E$ connections |
| $\alpha_1$ | -3.593 | appears in equation (29) |
| $\alpha_2$ | -14.16799 | appears in equation (29) |
| $\alpha_3$ | 1.7141 | appears in equation (29) |
| $\beta_1$ | 6.6686 | appears in equation (29) |
| $\beta_2$ | -2.9763 | appears in equation (29) |
| $\beta_3$ | 0.4271 | appears in equation (29) |
| $c$ | 0.3 | percentage of shared input from the total input |
| $\sigma$ | $\sqrt{25000}$ | standard deviation of the external Poisson process |

